# Uncoordinated long-patch base excision repair at juxtaposed DNA lesions generates a lethal accumulation of double-strand breaks

**DOI:** 10.1101/2020.11.15.383513

**Authors:** Kenji Shimada, Barbara van Loon, Christian B. Gerhold, Stephanie Bregenhorn, Verena Hurst, Gregory Roth, Cleo Tarashev, Christian Heinis, Josef Jiricny, Susan M. Gasser

## Abstract

Inhibition of the TOR pathway (TORC2, or Ypk1/2), or the depolymerization of actin filaments results in catastrophic fragmentation of the yeast genome upon exposure to low doses of the radiomimetic drug Zeocin. We find that the accumulation of double-strand breaks (DSB) is not due to altered DSB repair, but by the uncoordinated activity of base excision repair (BER) at Zeocin-modified DNA bases. We inhibit DSB formation by eliminating glycosylases and/or the endonucleases Apn1/2 and Rad1, implicating these conserved BER enzymes, or events downstream of them, in the conversion of base damage into DSBs. Among DNA polymerases, the reduction of Pol δ, and to a lesser extent Pol ε and Trf4 (a Pol β-like polymerase), reduces DSB formation. Finally, the BER enzymes, Ogg1 and AP endonuclease, are shown to co-precipitate with actin from yeast extracts and as purified proteins, suggesting that actin may interfere directly with the repair of Zeocin-induced damage.

## Introduction

Cells are exposed to various types of oxidative DNA damage arising from both endogenous metabolic activity and exogenous sources. Conserved DNA damage pathways guarantee genome integrity against these continual threats. Oxidative DNA base lesions are one the most frequent types of endogenous DNA damage, occurring at a very high rate (roughly 10^3^ events/cell/day) ^1^. Although base lesions are exploited in a controlled manner by cells undergoing differentiation or antibody diversification ^2, 3^, the spontaneous and random DNA base oxidation caused by respiration-generated Reactive Oxygen Species (ROS) is mutagenic if not properly repaired ^1^. Indeed, a common and highly mutagenic form of base oxidation, 7,8-dihydro-8-oxo-guanine (8-oxo-G), can trigger the C:G to A:T transversion mutations that are frequently found in lung, breast, ovarian, gastric and colorectal cancers ^4^. ROS are also generated by environmental factors, such as cigarette smoke and ionizing radiation (γIR).

Cells cope with DNA oxidation through a highly robust DNA repair pathway called base excision repair (BER) ^4, 5^. BER consists of a series of reactions generally initiated by DNA *N*-glycosylases. Monofunctional glycosylases recognize and remove damaged DNA bases, generating apurinic or apyrimidinic (AP) sites. Bifunctional *N*-glycosylases possess a DNA lyase activity as well, that cleaves the DNA backbone by means of a β-elimination reaction, generating a single-strand break (SSB) with a 3’-deoxyribosephosphate (3’-dRP) (reviewed in ^6^). Examples of bifunctional *N*-glycosylases are Ogg1, Ntg1 and Ntg2 in budding yeast (OGG1, NTH1 and NTH2 in mammals; ^7, 8^, although OGG1 possesses also monofunctional properties ^9^. AP sites and 3’-dRPs are then further processed by AP endonucleases (Apn1/APE1, Apn2/APE2) to produce single-strand breaks (SSB) terminating with a 3’-OH. This is a suitable substrate for polymerase-mediated elongation (reviewed in ^6^).

There are two BER pathways that repair Apn1/APE1-generated SSBs. The more common process in mammalian cells is Short-Patch BER (SP-BER), which uses DNA Polβ together with XRCC1 and DNA ligase III (*LIG3*). The second pathway, called Long-Patch BER (LP-BER), makes use of canonical replicative DNA polymerases δ and ε ^10, 11^, as well as DNA ligase I (Cdc9/LIG1), to fill-in the gap on the damaged strand and seal the remaining nick. Pol4, the yeast Polβ homologue, and Trf4 (Pap2) has been reported to have 2-deoxyribose-5-phosphate lyase activity, and may be implicated in SP-BER 12. However, there is no homologue of XRCC1 or ligase III in *S. cerevisiae*, leaving LP-BER as the dominant pathway in this organism ^13, 14^. LP-BER is also called Nucleotide incision repair, or NIR.

During BER, SSBs are generated by AP endonucleases as an intermediate in a multi-step process. SSBs, however, pose a serious risk to the genome, given that their presence on the template strand during replication or repair can generate a DNA double-strand break (DSB) ^15, 16^. To avoid this, gap-filling and sealing activities of BER must be coordinated with replicative and repair polymerases, particularly when two DNA base lesions are found in close proximity on opposite strands ^17^. How this avoidance or coordination is achieved is not well understood.

We previously showed that the regulation of actin polymerization by the TORC2 (target of rapamycin complex 2) pathway plays an important role in genome stability in budding yeast following exposure to either the radiomimetic antibiotic Zeocin or γIR, both of which generate strand breaks and oxidized base damage ^18, 19^. The combination of low-dose Zeocin with chemical inhibition of TORC2 provokes extensive chromosome fragmentation, with DSBs forming every 100-300 kb within 30 min (termed YCS for yeast chromosome shattering). TORC2 regulates the actin cytoskeleton in response to changes in plasma membrane status, through means of two effector kinases, Ypk1 and Ypk2 ^20^. Intriguingly, the perturbation of actin polymerization by Latrunculin A (LatA) appears to mimic TORC2 inhibition on Zeocin ^18^, suggesting that a loss of actin dynamics drives YCS (Hurst *et al*., in revision).

In recent years, several studies have implicated nuclear actin and actin-containing complexes in DNA repair, in some cases determined by the chromatin context of the damage. Whereas part of this may reflect a role for actin-containing chromatin remodeling complexes ^21, 22^, other roles have been proposed for actin itself in repair (reviewed in ^23 24^). One study in mammalian cells has argued that DNA damaging agents can induce actin filaments (F-actin) in the nucleus ^25^, while two others provide evidence that nuclear actin binding factors, namely WASP and Arp2/Arp3, contribute to DSB repair possibly by mediating repair focus formation ^26, 27^. Although nuclear F-actin can be detected only transiently, one cannot rule out a role in repair focus assembly and/or turnover, at least in higher eukaryotic cells.

In this study we have investigated the events that drive chromosome fragmentation after low dose exposure to the oxidizing agent Zeocin in combination with TORC2 inhibition. We show that the base excision repair machinery itself is responsible for YCS, and that along with AP endonucleases, gap-filling DNA polymerase activities (DNA Polδ) promote the formation of irreparable DSBs. Our data highlight the tight coordination that is necessary for the repair of oxidized DNA bases or abasic sites in close proximity, in order to avoid chromosome fragmentation. In another study (Hurst et al., in revision) we show that the toxic effect of TORC2 inhibition on AP endonucleases and LP-BER machinery is mediated by nuclear actin.

## Results

### TORC2 inhibition in combination with Zeocin provokes yeast chromosome shattering, unrelated to cell-cycle, HR, or NHEJ function

The related imidazoquinoline derivates NVP-BHS345 and CMB4563, both potent inhibitors of the yeast Tor2 kinase, cause rapid DSB formation and acute yeast chromosome fragmentation in combination with the radiomimetic drug Zeocin (Figure 1A) ^18^(Hurst *et al.*, in revision). Given the scale of DSBs in response to the low doses of Zeocin, the mechanism that underlies YCS is likely to result from a mis-regulated cellular pathway. Prime candidates for these are components of the DNA replication or repair machineries themselves. In an earlier study, the ablation of canonical DSB repair pathways, i.e. homologous recombination (HR) and non-homologous end-joining (NHEJ), did not mimic the YCS phenotype ^18^. However, given recent evidence implicating nuclear actin in homology-dependent DNA repair in human and *Drosophila* cells ^26, 27^, we have examined the type of DNA damage involved in chromosome shattering and have quantified our gels to be able to compare a range of conditions (see Material and Methods).

**Figure 1.**
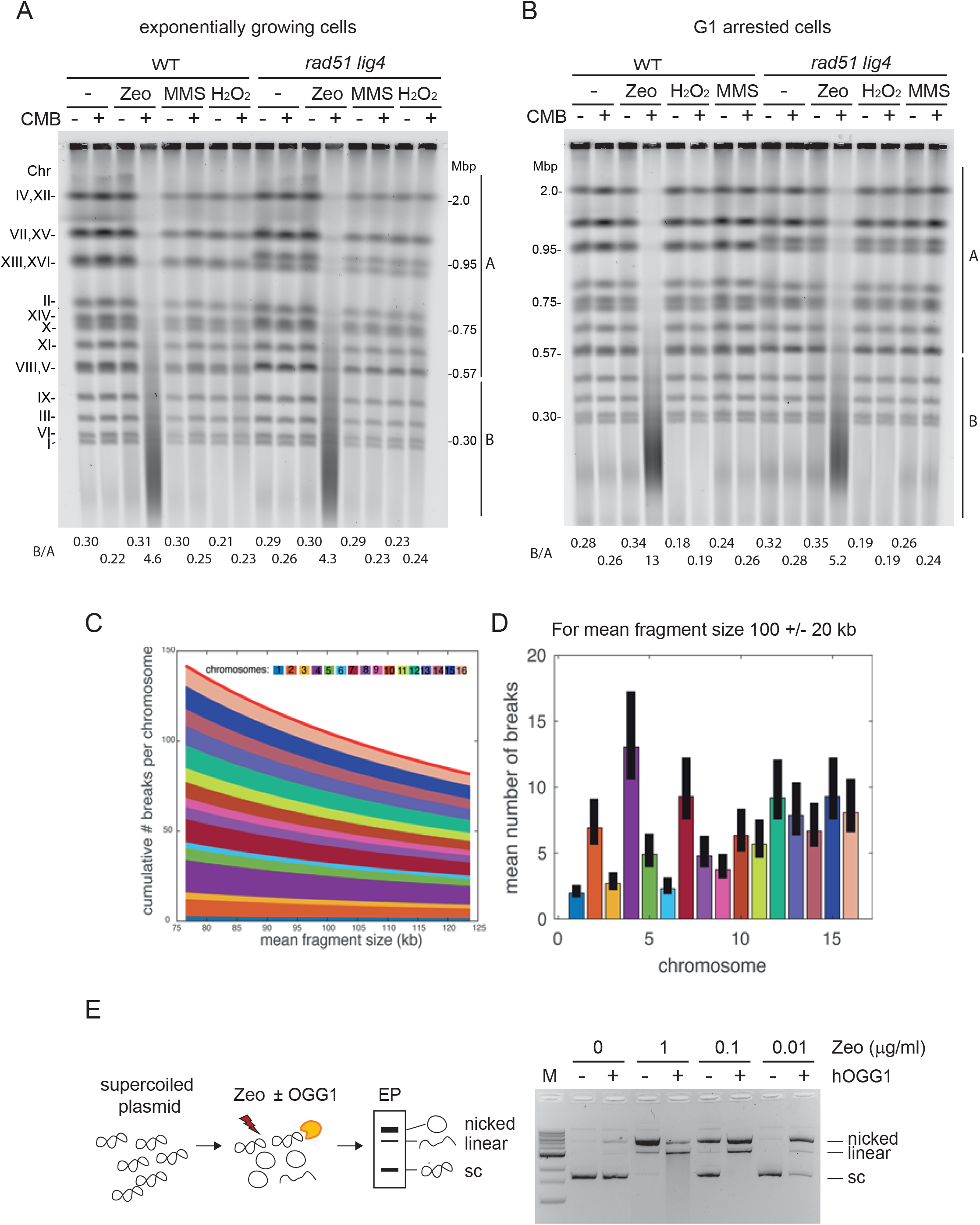
TORC2 inhibition induces YCS at sites of Zeocin-generated DNA base oxidation, independent of S phase and DSB repair pathways. A, B) Massive chromosome fragmentation occurs irrespective of DSB repair and independent of cell cycle. Wild-type (WT, GA-1981) and an isogenic *rad51 lig4* double mutant (GA-6098) were grown exponentially (A), or were arrested in G1 by α-factor for 1.5h (B). Cells were treated with 50 μg/ml Zeocin, 1 mM H_2_O_2_, or 0.05% MMS alone, or in combination with 0.5 μM CMB4563 (a potent TORC inhibitor) in yeast^31^ (Hurst *et al*., in revision) for 80 min. Genomic DNA was isolated in the agarose plug at 30°C as described in Material and Methods, then analysed by CHEF gel analysis and stained with SYBR safe to visualize genomic DNA. Signal intensity of each lane was measured by Image J program then the signal ratio of B (below 0.57 Mbp till ~20 kbp) over A (covering over the largest chromosome band 4 and 12 till chromosome 5 and 8 band) is indicated below the gel image. C) For a given mean fragment size, we calculate the mean number of breaks for each chromosome and pile those numbers from chromosome 1 at the bottom to chromosome 16 at the top. The solid red line represents the mean number of breaks in the genome. D) Theoretical mean number of breaks per chromosome for a mean fragment size of 100kb. The solid black bars represent the values for a 20% error in the estimated mean fragment size (i.e. the real mean fragment size is between 80kb and 120kbD. E) Zeocin treatment induces substrate for hOGG1 on supercoiled plasmids *in vitro*. Supercoiled plasmid DNA was mixed with the indicated concentration of Zeocin for 60 min at 37°C, and then further incubated with or without purified hOGG1 for 30 min at 37°C. Plasmid DNA was purified and loaded on an agarose gel. Positions of nicked, linearized and supercoiled plasmids are indicated besides the gel image.

We challenged yeast cells with three different DNA damaging reagents independently: Zeocin, H_2_O_2_, and MMS, and compared the integrity of the 16 yeast chromosomes of a wild-type strain and a double mutant defective both for HR (*rad51*) and NHEJ (*lig4*), by CHEF gel analysis. Zeocin is a radiomimetic agent from the bleomycin family of antibiotics that causes ss and ds breaks, base oxidations and AP sites, that are often found closely juxtaposed on opposite strands of the DNA ^19, 28^. H_2_O_2_ and MMS cause random DNA oxidation and alkylation of DNA bases, respectively. Several forms of alkylated DNA bases block replicative polymerases, and thus MMS treatment also induces a strong replication fork-associated damage (reviewed in ^29^). We carefully avoided heat-labile MMS-induced lesions by preparing yeast DNA at 30°C ^30^, and found no chromosome fragmentation after treatment with MMS nor with H_2_O_2_, either alone or together with TORC1/2 inhibition ^31^ (Figure 1A). In contrast, the combination of Zeocin with CMB4563 led to full YCS (Figure 1A). This was equally efficient in the absence of the two DSB repair pathways (*rad51 lig4*), confirming that YCS does not reflect inhibition of the canonical DSB repair pathway, nor does general alkylation or oxidation sensitize cells to the TORC2 inhibitor (Figure 1A).

Because DSB repair mechanisms show strong cell cycle preferences, i.e., G1>S for NHEJ and S>G1 for HR, we also examined the effect of G1 arrest on the combination of damage and DSB repair pathway mutants. Again, only the combination of Zeocin and CMB4563 triggered YCS, and the result was identical in the presence or absence of functional HR/NHEJ repair pathways (Figure 1B). Thus, not only is chromosome fragmentation independent of the DSB repair machinery, but it also does not depend on an S-phase specific mechanism (Figure 1B).

We quantified the number of DSBs incurred using our protocol to yield an average fragment size of 100 ± 20 kb based on a random introduction of DSBs in the yeast genome (1.2 × 10^7^ bp). We calculate between 80 and 140 DSBs per genome (Figure 1 C,D). Given that we see very little chromosome fragmentation in the presence of Zeocin without the imidazoquinoline, we infer that the breaks are avoided by means of another repair pathway.

A unique feature of the reagent Zeocin is that it attacks the DNA backbone at closely juxtaposed sites, often generating two nicks in close proximity^19^. One model suggests that it does this by producing a hydroxyl radical that would also cause DNA base oxidation ^28, 32^. Two lesions immediately adjacent to each other could thus generate a DSB, if processed by a nicking enzyme. We therefore checked whether Zeocin can induce oxidized 7,8-dihydro-8-oxo-guanine (8-oxo-G) *in vitro,* which could then serve as an efficient substrate for OGG1. OGG1 is a bifunctional N-glycosylase that generates an abasic site at 8-oxo-G, and then which can cleave the DNA backbone by means of a β-elimination reaction through its DNA lyase activity^33^. We examined the effects of Zeocin-induced damage, and its susceptibility to OGG1 action, by incubating a closed circular DNA plasmid with the drug, with or without purified human OGG1. In the absence of OGG1, high doses of Zeocin alone (0.1-1 μg/ml) primarily generated nicked DNA fragments, with only 50% efficiency at 0.1 μg/ml Zeocin (Figure 1E). In the presence of OGG1 at this concentration Zeocin, all supercoiled plasmid was converted either to a nicked or a linearized form, reflecting single- and double-strand scission, respectively (Figure 1E). Even, at ten-fold less Zeocin (0.01 μg/ml) with OGG1 present, nearly all plasmid template was nicked. Thus closely juxtaposed Zeocin-induced lesions can generate DSBs *in vitro*, particularly if processed by OGG1. Given that OGG1 is the first enzyme in the BER multi-step pathway (Figure 2A), we explored further the role of BER enzymes in YCS.

**Figure 2.**
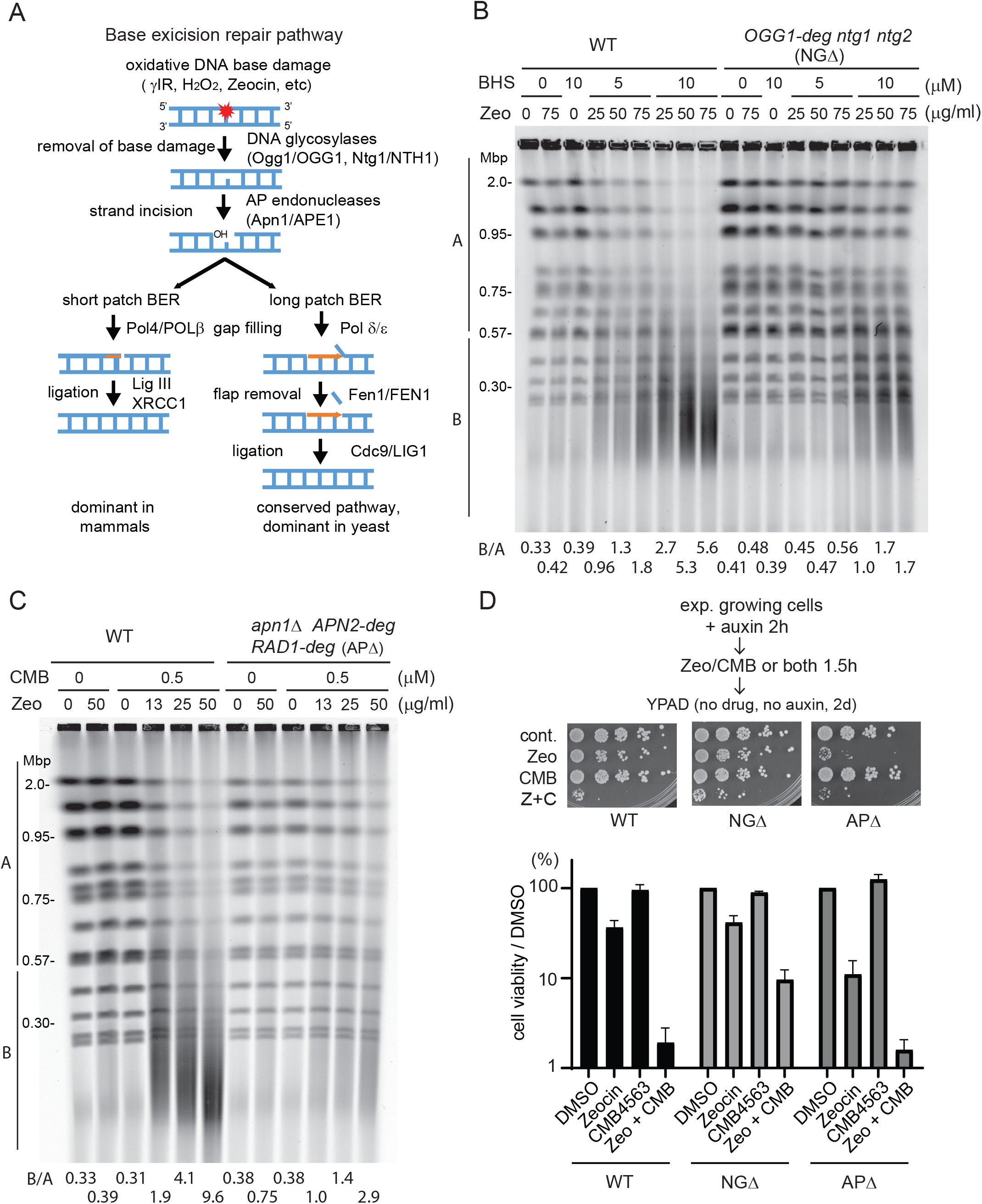
BER *N*-glycosylase enzymes and AP nuclease activities are responsible for YCS. A) Scheme of SP- and LP-BER pathways with yeast enzymes in small letters and mammalian enzymes in all capitals. B) Loss of *N*-glycosylase activity confers resistance to YCS. Exponentially growing wild-type (GA-8369) and *Ogg1-deg ntg1 ntg2* (GA-8457, NGΔ) cells were treated with 0.5 mM IAA (indole acetic acid) for 4h. Cells were then treated with DMSO control, NVS-BHS867^18^ (TORC inhibitor; Hurst *et al*., in revision), Zeocin, or a combination of both drugs as indicated for 70 min. Genomic DNA was analysed by CHEF gel analysis and visualized using SYBR safe. C) Loss of AP endonuclease activity strongly inhibits YCS. Exponentially growing wild-type (GA-8369) and *apn1 APN2-deg RAD1-deg* (GA-8509, APΔ) cells were treated with 0.5 mM IAA for 1h to deplete Apn2 and Rad1. Cells were then incubated with 0.5 μM CMB4563 in the presence of Zeocin as indicated for 70 min. Genomic DNA was revealed by CHEF gel analysis. D) Wild-type, NGΔ, and APΔ cells were exponentially grown in complete minimal media (SC), then 0.5 mM IAA was added for 2h. Cells were then treated with 50 μg/ml Zeocin, 0.5 μM CMB4563, or a combination of both drugs for 90 min. Cells were spotted on YPAD in 5-fold dilution series. A picture was taken after 2 days and the percentage cell viability was plotted, normalized by DMSO control. Error bar represents standard error of 5 biological replicates.

### BER glycosylases and AP nucleases are implicated in YCS

To test if and how BER activity contributes to chromosome shattering *in vivo*, we generated a yeast strain in which all three *N*-glycosylases that initiate oxidative DNA base repair can be inactivated (Figure 2A; *OGG1-*auxin dependent degron *ntg1Δ ntg2Δ;* NGΔ). We cultured NGΔ and the corresponding wild-type cells exponentially in the absence of auxin, and then added 0.5 mM Indole-3-acetic acid (IAA) to the culture for 4h to deplete Ogg1 prior to drug treatment. Cells were subsequently treated with Zeocin, yeast TOR inhibitor NVP-BHS345, or a combination of the two at different concentrations, as indicated (Figure 2B). In wild-type cells, we observed chromosome shattering with the combined treatment of Zeocin and NVP-BHS345, as expected (Figure 2B). However, by reducing *N*-glycosylase activity we significantly reduced chromosome shattering (Figure 2B), suggesting that DSB formation depends on the BER pathway. Loss of Ogg1 alone, as a full *ogg1* deletion, was not sufficient to confer strong YCS resistance, arguing that the repair factors Ogg1, Ntg1 and Ntg2 are redundantly responsible for the DSBs that occur during YCS (Suppl. Figure 1A).

One of the key downstream effects of TORC2 activation is the control of cytoplasmic actin polymerization. We therefore asked whether the loss of the three *N*-glycosylases also conferred resistance to chromosome fragmentation, not only induced by Zeocin and TORC2 inhibition, but also by Latrunculin A (LatA), which similarly depolymerizes actin. Indeed, the induction of YCS by the combination of Zeocin and LatA was strongly reduced in NGΔ cells, as observed following treatment with Zeocin and CMB4563 (Suppl. Figure 1B).

### Loss of AP endonuclease activity results in a strong resistance to YCS

We next asked if AP endonucleases, which act downstream of the *N*-glycosylases to cleave the DNA backbone, were implicated in YCS. Yeast Apn1 and Apn2 possess canonical AP endonuclease activity, while Rad1-Rad10 was shown to have a compensatory function in their absence. All have been implicated in BER, usually functioning at AP sites generated by *N*-glycosylases ^34^. We generated an *apn1Δ APN2-degron RAD1-degron* strain (APΔ) to eliminate all AP endonuclease activity.

Exponentially grown wild-type and APΔ cells were incubated with 0.5 mM IAA, followed by a treatment with Zeocin, CMB4563, or a combination of both compounds. Strikingly, the triple knock-out APΔ cells were almost completely resistant to YCS triggered by the TORC2 inhibition (Figure 2C). Once again this illustrated that the BER pathway itself contributes to the generation of DSBs under these conditions.

The *apn1 apn2* double mutant exhibited incomplete resistance to YCS, confirming that Rad1-Rad10 can contribute to the toxic processing events (Suppl. Figure 1C). Importantly, we could partially suppress YCS by expressing *APN1* from a single-copy plasmid (pAPN1) in APΔ cells. This indicates again that Apn1, Apn2 and Rad1/10 have partially redundant activities, each able to contribute to YCS on low-level Zeocin and TORC2 inhibitor (Suppl. Figure 1D). This control also excludes the possibility of a background mutation in the APΔ cells that might suppress the shattering phenotype.

We asked if the resistance to chromosome shattering observed in the APΔ and NGΔ strains correlated with improved colony survival after transient Zeocin and CMB4563 treatment. First, we note that in wild-type cells only a small fraction of cells survive the combined Zeocin and CMB4563 treatment, whereas the survival of either treatment alone is extremely efficient (Figure 2D). In the APΔ strain, cell viability dropped to roughly 10% after treatment with Zeocin alone, and the combined treatment yielded the same low survival as wild-type cells, despite their resistance to chromosome shattering (Figure 2C, 2D). These data underscored the essential role that AP endonucleases play in the survival of Zeocin-induced damage and illustrated that resistance to chromosome shattering does not necessarily correlate with improved cell viability. In brief, both *N*-glycosylases and AP nucleases are implicated in DSB generation arising from base oxidation, as well as being essential for recovery from Zeocin-induced damage. Intriguingly, the loss of *N*-glycosylases actually improved cell survival over that of wild-type cells following the combined CMB4563 + Zeocin treatment (Figure 2D), suggesting that in the absence of base removal, an alternative pathway of repair may take place. In conclusion, these results argue that AP endonucleases and/or the downstream BER events are responsible for the DSBs that drive YCS.

### Loss of Pol32, a subunit of DNA Polδ, reduces the chromosome fragmentation

As described above, AP endonuclease activity produces a single strand break with a 3’-OH and 5-deoxyribosephosphate (5-dRP), which is subsequently processed either by SP- or LP-BER ^4^. Several SP-BER factors, namely DNA ligase III and XRCC1, are missing in *S. cerevisiae*, therefore yeast depends on the replicative DNA Polδ and Polε, along with DNA ligase 1 in LP-BER ^14, 35^. DNA Polδ (Pol3) is known to preferentially fill-in short single-stranded gaps, serving as a key repair polymerase in addition to its essential function in lagging strand synthesis ^36^. Besides a role in BER, the nonessential Polδ regulatory subunit Pol32 appears to help bypass of AP sites ^37^. Yeast also encodes a mammalian DNA Polβ homologue, Pol4, that was implicated in NHEJ ^38, 39^. Given their potential roles in BER, we tested strains lacking the genes encoding Pol4 and/or Pol32 for the YCS phenotype.

Whereas the *pol4* deletion had no effect, the *pol32* null allele resulted in reduced chromosome fragmentation (Figure 3A), and the *pol4 pol32* double mutant showed the same level of resistance to YCS as the *pol32* single mutant (Figure 3A). This implicates Pol32 but not Pol4 in generating YCS. The trans-lesion synthesis polymerase Polζ shares the Pol32 subunit with Polδ ^40^, so we also tested whether deletion of *REV3*, a catalytic subunit of Polζ, confers resistance to YCS. However, the level of chromosome shattering in *rev3* cells was identical to that in wild-type cells (Suppl. Figure 2A), suggesting that Polδ, rather than Polζ, promotes DSB formation following Zeocin and CMB4563 treatments.

**Figure 3.**
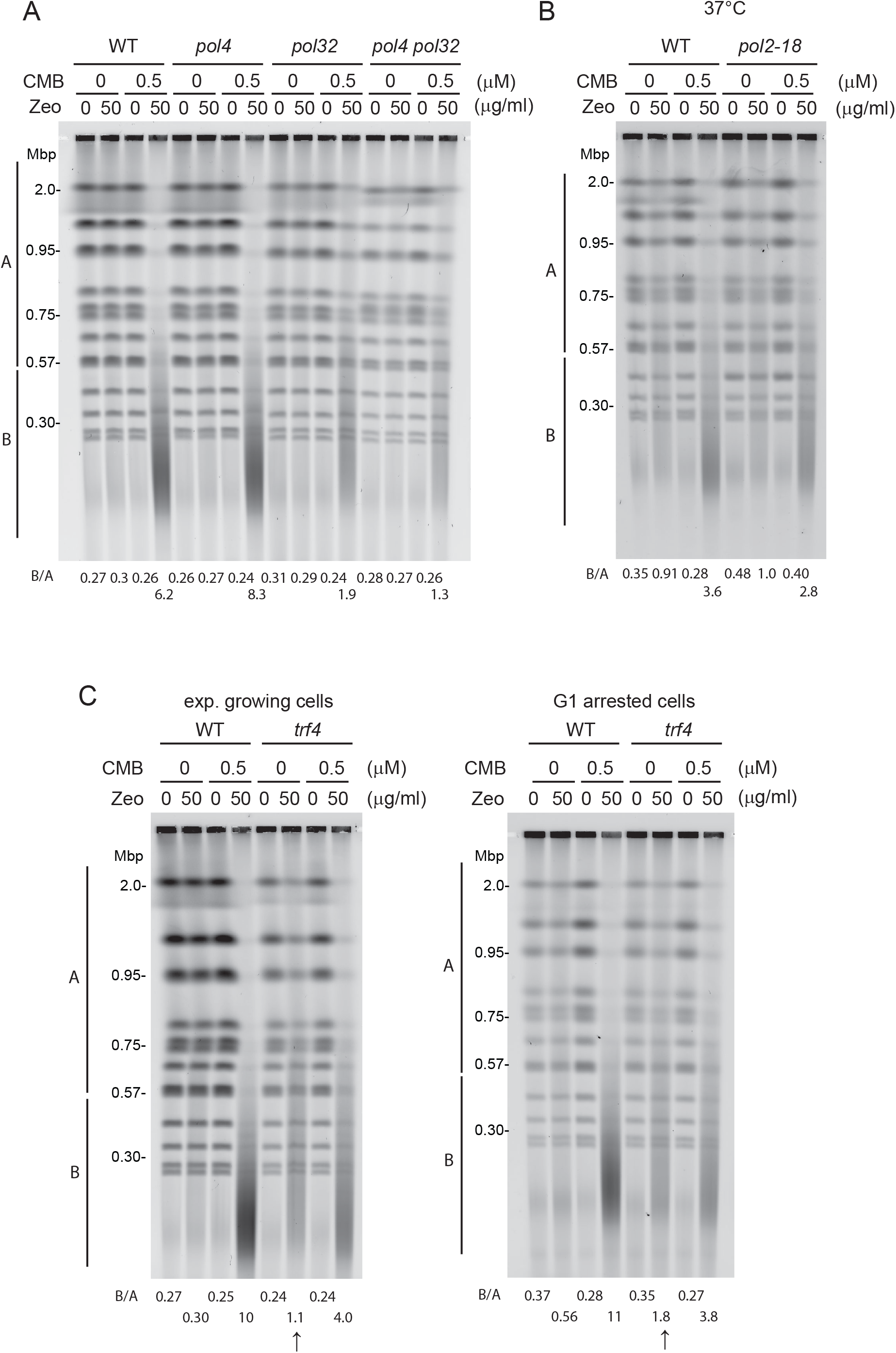
Impaired gap filling by DNA Polδ and Polε alter YCS efficiency. A) DNA Polδ activity facilitates YCS. Wild-type (GA-1981), *pol4* (GA-10595), *pol32* (GA-9686), and *pol4 pol32* (GA-10697) mutants were exponentially cultured in SC medium. Cells were treated with 0.5 μM CMB4563, 50 μg/ml Zeocin, or a combination of both reagents for 60 min. Genomic DNA was subjected to CHEF gel analysis. B) Loss of DNA Polε activity affects YCS. Wild-type (GA-741) and *pol2-18* (GA-742) cells were exponentially cultured in SC at 25°C. Cell culture was shifted to 37°C for 40 min, then treated with 1 μM CMB4563, 75 μg/ml Zeocin, or a combination of both reagents for 70 min, prior to CHEF gel analysis. C) Loss of Trf4 enhances Zeocin sensitivity but reduces YCS. Wild-type (GA-1981) and *trf4* (GA-10632) cells were exponentially cultured in SC, then treated with 50 μg/ml Zeocin, 0.5 μM CMB4563, or a combination of both drugs for 80 min, prior to CHEF gel analysis (left hand gel). The same procedure was performed on cells synchronized and arrested in G1 phase by a pre-incubation with α-factor. All subsequent steps were identical.

It has been proposed that DNA Polε functions redundantly or alongside Polδ in BER ^10, 41^. Another study showed that Trf4 possesses dRP lyase activity relevant to Polβ function ^12^. Therefore, we tested both the effect of *trf4* deletion and of the *pol2-18* allele, a temperature-sensitive mutation in the catalytic subunit of Polε, for a role in YCS. We find that the *pol2-18* mutation conferred a very minor resistance to YCS at 37°C, (Figure 3B), while the *trf4* mutant showed hypersensitivity to Zeocin alone, leading to limited DNA fragmentation in the absence of CMB4563. This was true both in exponential and G1-arrested cultures (Figure 3C). In contrast, under YCS conditions the loss of Trf4 also conferred a minor resistance to chromosome fragmentation (CMB + Zeo, Figure 3C). Taken together, our results indicate that the gap-filling activity in LP-BER of juxtaposed lesions is primarily fulfilled by DNA Polδ, although there may be minor contributions from DNA Polε and the non-canonical BER enzyme Trf4, which acts by removal of the 5’-terminal abasic residue generated by AP endonuclease incision (like mammalian pol β) thus contributing to SP-BER. These observations would also be consistent with recent observations showing that DNA Polε also can contribute to SP-BER ^42^.

A further pathway relevant to base oxidation is the incorporation of free oxidized nucleotides during DNA replication and repair, which has been reported to result in genome instability and significant cytotoxicity ^43, 44^. Loss of hMTH1, which removes free oxidized purine nucleotides ^45^, sensitizes the cell to oxidative damage in a manner partly dependent on Polβ ^44^. We hypothesized that the observed increase in chromosome fragmentation, insofar as it is dependent on DNA Polδ and Polε, might reflect the incorporation of oxidized free nucleotides at sites of Zeocin-induced damage. Therefore, we examined if YCS would be enhanced by an increase in free oxidized nucleotides, which we triggered by deleting *PCD1*, the yeast homologue of hMTH1. We found that YCS was almost identical in wild-type and *pcd1Δ* cells (Suppl. Figure 2B). Moreover, the overexpression of Pcd1, done in an attempt to sanitize oxidized purine nucleoside triphosphates, did not reduce the level of YCS (data not shown). These results argue that the incorporation of oxidized nucleotides (e.g., 8-oxo-G) by DNA polymerases is not the driver of YCS. Rather chromosomal DSBs arising from Zeocin treatment stem from the processing of oxidized bases within genomic DNA.

### AP endonucleases, but not *N*-glycosylases, play key roles in YCS in G1 phase cells

Since YCS can be induced in G1 phase in the absence of ongoing DNA replication (Figure 1A; ^18^, we next tested the role of the BER factors on YCS in α-factor-arrested G1-phase cells. Exponentially growing wild-type and NGΔ cells were arrested in G1 using α-factor and were tested for YCS. Surprisingly, we observed little or no YCS resistance in G1-arrested cells with depleted *N*-glycosylase activity compared to exponentially growing cultured conditions (Figure 4A, cf. Figure 2B). Even when random and synchronized cells were tested directly in parallel, NGΔ cells were resistant to YCS in exponential growth, but sensitive when G1-arrested (Suppl. Figure 3A). In contrast, the loss of AP endonuclease activity, still conferred a strong resistance to YCS in G1 (Figure 4B), suggesting that AP endonuclease or a repair activity downstream from AP enzymes drives chromosome fragmentation at all cell cycle stages.

**Figure 4.**
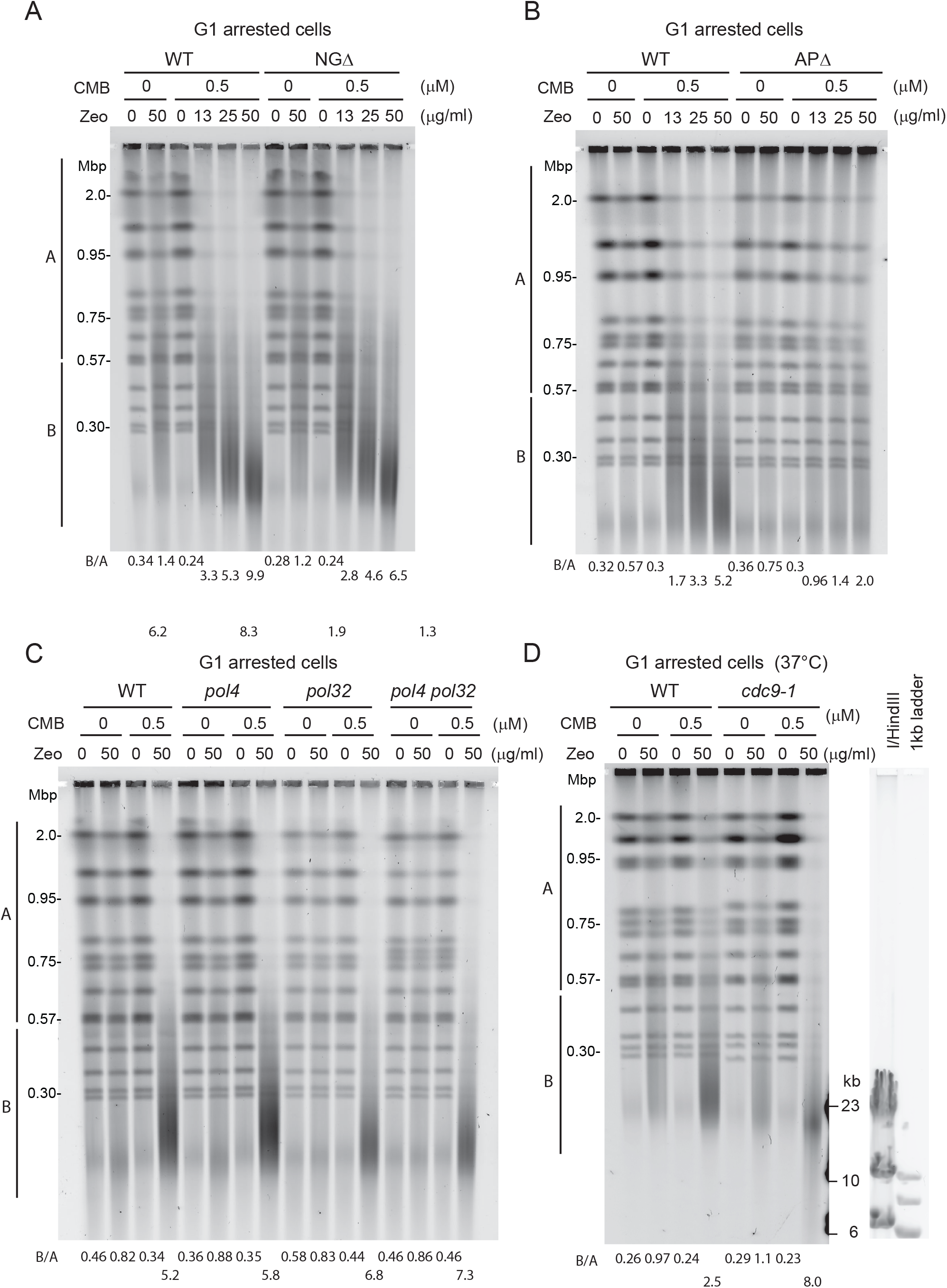
AP endonuclease activities, but not *N*-glycosylases, are is required for YCS in G1 cells. A) *N*-glycosylase activity is largely dispensable for YCS in G1-arrested cells. Exponentially growing wild-type (GA-8369) and *Ogg1-deg ntg1 ntg2* (GA-8457) cells were treated with 0.5 mM IAA for 2h, then α-factor was added to the culture to arrest in G1 for another 2h. Cells were then treated with DMSO control or CMB4563 in the presence of different amounts of Zeocin as indicated for 80 min. Genomic DNA was revealed by CHEF gel analysis. B) YCS occurs in an AP-endonuclease-dependent manner in G1. Exponentially growing wild-type (GA-8369) and *apn1 APN2-deg RAD1-deg* (GA-8509) cells were treated with 0.5 mM IAA and α-factor for 2h. Cells were treated with CMB4563 and Zeocin as indicated for 80 min, followed by CHEF gel analysis. C) Loss of Pol32 does not reduce YCS efficiency in G1-arrested cells. Exponentially growing wild-type (GA-1981), *pol4* (GA-10595), *pol32* (GA-GA-9686), and *pol4 pol32* (GA-10697) mutant cells were arrested in G1 by α-factor for 100 min. Cells were then treated with 0.5 μM CMB4563, 50 μg/ml Zeocin, or a combination of both reagents for 70 min prior to CHEF gel analysis. D) Loss of DNA ligase I (Cdc9) activity greatly enhances YCS. Wild-type (GA-8709) and *cdc9-1* (GA-8708) cells grew exponentially in SC at 25°C. Cells were arrested in G1 by α-factor for 2h, then shifted to 37°C for 45 min. Cells were treated with 0.5 μM CMB4563, 50 μg/ml Zeocin, or a combination of both reagents for 60 min prior to CHEF gel analysis.

We further examined the contributions of DNA polymerase activities in G1-arrested cells. Interestingly, neither the *pol4* the *pol32* nor the double mutant had any effect on YCS in G1 phase, unlike their impact on exponentially growing cells (compare Figures 3A and 4C). Nor did we detect an impact of the *pol2-18* mutation on YCS, while *trf4* deletion conferred a slight resistance to YCS in both G1 and random cultures (Figure 3C). This could arise either from additional redundancy among polymerases in G1 phase, or from gaps in G1 phase being smaller and requiring less DNA synthesis.

Once a DNA polymerase has filled the gap left by base excision, ligases must act to avoid a persistent nick. If two nicks are near each other, on opposite strands, failed re-ligation could also lead to DSBs. We therefore tested whether the loss or reduction of DNA ligase activity would generate DSBs in the presence of Zeocin, enhancing YCS. We used the temperature-sensitive ligase I allele *cdc9-1,* and tested its effect on YCS in G1 phase, thus avoiding the high level of replication intermediates inherent to this mutant, which would interfere with CHEF gel analysis. Indeed, inactivating Cdc9 (Figure 4D), but not ligase 4 (Figure 1A, B), strongly enhanced YCS; moreover, it generated fragmentation on Zeocin, even in the absence of TORC2 inhibition. The size of the fragments produced by YCS were ~ 20 kb, that is, even smaller than the usual ~100kb YCS-generated fragments.

This suggests that Lig1 is repairing Zeocin-induced damage even under YCS conditions; its loss generates roughly twice as many DSBs as in wild-type cells (1 per 20kb, rather than 1 per 50-100kb). Intriguingly, yeast Cdc9/Lig1 is one of the factors that is phosphorylated under YCS conditions (Hurst *et al.*, in revision), suggesting that it may be upregulated to cope with excessive damage. Importantly, the absence of Lig1, unlike the loss AP endonucleases, accentuates rather than reduces YCS.

Our data suggest that AP endonuclease activity contributes to YCS in G1 phase, surprisingly without prior action of a BER *N*-glycosylase. We note that G1-arrested cells exhibit a much higher sensitivity to 50 μg/ml Zeocin than cycling cells, resulting in only 1% viability even in a wild-type background (Suppl. Fig 3B). One explanation for this may be that Zeocin is more potent in G1, generating AP sites that bypass the need for *N*-glycosylase. However, we did not detect more AP sites after Zeocin treatment in α-factor arrested cultures than we did in those in exponential growth (Suppl. Figure 3C). This was true either with or without Zeocin, CMB4563 or the combined treatment. MMS, on the other hand, was previously reported to generate a very high number of AP sites through the action of the Mag1 *N*-glycosylase ^46, 47^.

A hyper-radical reagent often generates a “dirty end”, that can inhibit the action of repair polymerases and ligases. Since mammalian Polβ can remove 2-deoxyribose-5-phosphate (5-dRP) or 5’-AMP-dRP from the 5’ termini of BER intermediates, we asked if other related enzymes might be contributing to YCS in G1 phase in yeast. To examine this, we deleted *tpp1*, which encodes a protein that removes 3’ phosphates at strand breaks, and *hnt3*, which encodes aprataxin, an enzyme that reverses DNA adenylation (5’-AMP-dRP), as well as *tdp1*, which encodes an enzyme that hydrolyses 3’ and 5’ phosphotyrosyl bonds, such as those created by cleavage by topoisomerases I or II, respectively ^48, 49, 50^. However, we found no YCS resistance of *tpp1, hnt3* nor *tdp1* deletion strains (Suppl. Figure 4A,B), and only detected slight sensitivity to Zeocin in the absence of TORC2 inhibition, with low level fragmentation. This may indicate that the enzymes do process blocked ends to promote repair in G1-phase cells, perhaps acting upstream of Lig1, promoting the repair of a subset of Zeocin-induced lesions. However, unlike AP enzymes, their absence does not confer resistance to YCS, thus their activities are not required to generate DSBs under YCS-inducing conditions.

Our data clearly suggest that AP endonucleases, which act independently of glycosylases in G1 phase, are the key enzymes whose activity (or misregulation) drives chromosome fragmentation under YCS conditions. A previous report had implicated Apn1 in an Ogg1-independent repair of 8-oxo-G residues thanks to its progressive 3’ to 5’ exonuclease activity ^51^. In this so-called nucleotide incision repair (NIR) pathway, Apn1’s nuclease activity acts directly at modified bases to generate a 3’ OH end ^52, 53^. Thus, we speculate that Apn1 activity may contribute to YCS in G1-arrested cells through NIR. Consistent with this hypothesis, elevated levels of Apn1 or Apn2 provokes strong Zeocin sensitivity in yeast, particularly when coupled with a nuclear localization signal (NLS) (Suppl. Figure 4C).

### BER factors interact with F-actin

Our previous results indicated that the usually robust BER pathway is somehow subverted by cytoplasmic F-actin depolymerization to generate breaks^18^ (Hurst *et al.*, in revision). The stability of the actin cytoskeleton itself is regulated by TORC2 and a subsequent cascade of kinases including Ypk1/2, Frk1, and Ark1/Prk1 and AKl1 ^20^. The effect can be mimicked by LatA treatment or by the overexpression of actin lacking a functional nuclear export signal, which leads to its accumulation in the nucleus (Hurst *et al*., in revision).

Given the links between actin dynamics and YCS, we hypothesized that actin might bind BER factors. Due to the paucity of anti-actin antibodies and the deleterious effect of adding epitope tags to actin ^54, 55^, we made use of novel G- or F-actin specific bicyclic peptides isolated from a phage display library ^56^. These bicyclic peptides are circularized sequences of 10-17 amino acids with highly selective binding specificity for different forms of actin, validated with both *in vitro* and *in vivo* assays ^56^. We coupled the peptides A18 (F-actin specific), A15 (F-/G-actin), or the TATA2 control peptide to magnetic beads, and recovered their ligands and associated factors from a total yeast cell extract, after a 2h incubation and rigorous washing of the recovered complexes (see Materials and Methods). The proteins recovered were visualized by silver staining and the more prominent peptides were identified by mass spectrometry (Figure 5A). We found that A18 and A15 efficiently bind actin, as expected, and along with actin we recovered many abundant and well-characterized actin-binding proteins from the cell extract, including Pan1, Sla2, Las17 and Cofilin, along with two subunits of the ssDNA binding factor, RFA. Using Western blots to probe the bicyclic peptide-recovered factors, we found a strong enrichment for Apn1 in both the A18 and A15 pull-down fractions, which was not the case for another nuclear factor (Mcm2), nor for tubulin (Figure 5A). Ogg1 was weakly detected in the A15 pull-down only.

**Figure 5.**
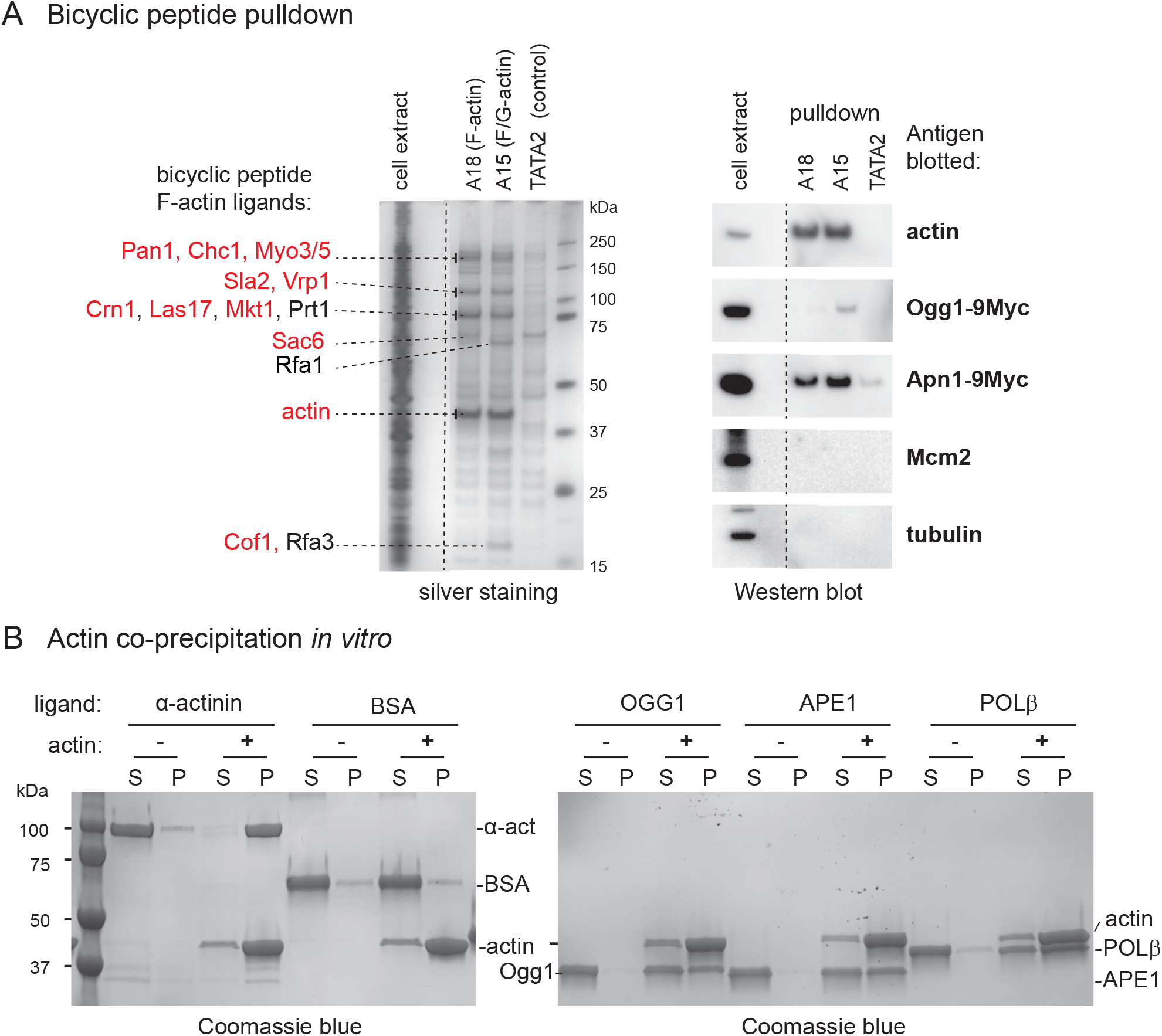
BER factors bind actin in yeast extract and *in vitro*. A) Apn1 and Ogg1 co-precipitate with F-actin from yeast cell extract. Bicyclic peptides specific for F-actin (A18), or F/G actin (A15), or a linker peptide TATA2 were linked to magnetic beads and incubated with total yeast cell extract. The proteins recovered are visualized by silver staining and prominent proteins identified by mass spectrometry are indicated. The same fractions were probed by Western blots for the indicated factors: Apn1, Ogg1, Mcm2 and tubulin. B) Purified BER factors (human OGG1, APE1, or POLβ) were incubated with purified rabbit actin. α-actinin and BSA were used as positive and negative controls, respectively. After incubation at room temperature for 30 min, proteins were recovered by centrifugation at 250.000 × g for 30 minutes. Proteins sedimenting with actin were denatured and analysed by gel electrophoresis and Coomassie Blue staining

We next asked if the interaction between BER factors and F-actin was direct, and whether the interaction could be extended to homologues in other species. We incubated highly-purified human OGG1, APE1, or POLβ and purified rabbit actin, and tested for interaction *in vitro*. The known actin ligand α-actinin served as a positive control and BSA as a control for nonspecific interaction. Because actin polymerizes spontaneously *in vitro*, we could recover it and its associated proteins by centrifugation. The α-actinin co-precipitated efficiently with F-actin under these conditions, while BSA did not (Figure 5B). Interestingly, a significant fraction (40-50%) of OGG1, APE1, POLβ were also recovered in the F-actin pellet, while they did not sediment under identical conditions when actin was omitted from the reaction (Figure 5B). This provides evidence that purified BER factors can interact with actin directly and suggests a means through which TORC2 inhibition and LatA-driven disassembly of the actin cytoskeleton might regulate BER.

## Discussion

Earlier work demonstrated a rapid and extensive fragmentation of the yeast genome following the exposure of wild-type cells to low doses of Zeocin and non-toxic levels of a TORC2 inhibitor, neither of which alone affected genomic integrity^18^. In Hurst *et al*. (in revision), we show that actin depolymerization is at the root of the TORC2 inhibitor effect. Here we describe the molecular events that lead to a synergistic collapse of genomic integrity, following exposure to Zeocin. The observed genomic fragmentation requires enzymes that are central for the processing of oxidized base residues during BER, notably *N*–glycosylases and AP nucleases. Oxidized bases such as 8-oxo-G occur frequently in the genome, and are generally repaired throughout the cell cycle by either short-patch or long-patch BER (reviewed in ^4^). A key characteristic of Zeocin, and related compounds like bleomycin, is their propensity to induce closely-positioned oxidative damage on paired nucleic acid chains ^19^. This is not the case for ionizing radiation (γIR), H_2_O_2_, nor for other commonly used alkylating or base-oxidizing agents, which do not trigger YCS as acutely. The closely juxtaposed Zeocin-induced lesions must be processed by *N*-glycosylases and AP endonucleases in a sequential manner to allow repair of two adjacent modified bases without generating a DSB (see Figure 6).

**Figure 6.**
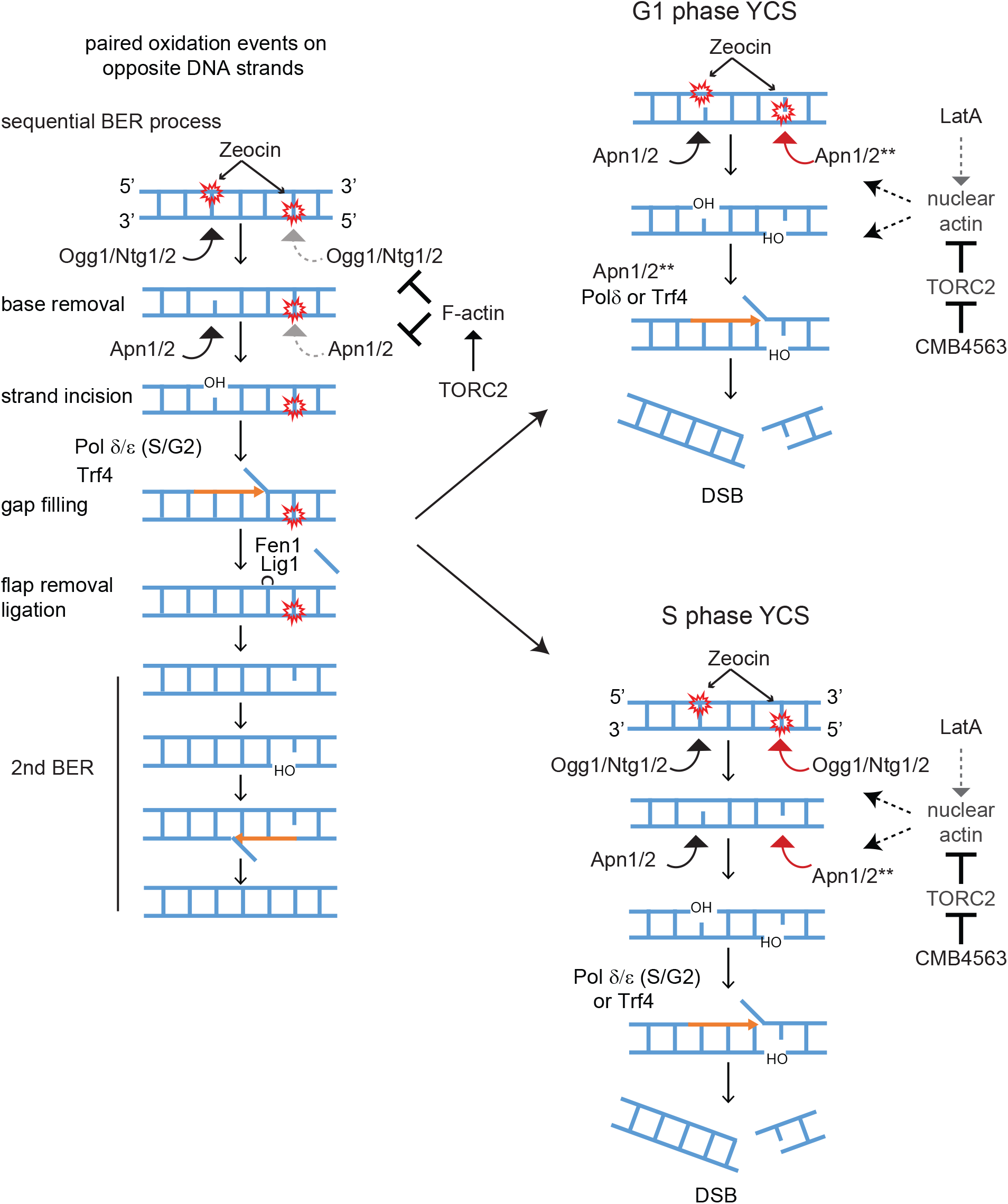
Model of correct and actin-misregulated BER of adjacent lesions. We propose that the repair of closely juxtaposed oxidized bases on DNA becomes toxic in the presence of reagents that depolymerize actin in the cytoplasm, due to a miscoordination of the two step BER process. The undisturbed sequential repair of two closely positioned oxidation events is shown on the left. The activities of Ogg1 and Apn1 may be indirectly kept in check by cytoplasmic F-actin. In G1 phase cells the increase in nuclear actin may activate Apn1/2 (see **) to drive premature processing of both lesions. In S phase the impact of nuclear appears to be both at the level of the glycosylases as well as AP endonucleases. We do not rule out that the unconstrained elongation by DNA Polδ or Trf4, also contribute to DSB formation. For further details see Discussion.

We find that F-actin depolymerization allows Zeocin-induced base lesions to be rapidly converted into irreparable DSBs, in a manner dependent on key enzymes of the BER pathway. In S and G2 phase cells, the loss of *N*-glycosylases or of the AP endonucleases, both prevent generation of DSBs, but in pheromone-arrested G1 phase cells, the loss of *N*-glycosylases does not block YCS, while ablation of the nucleases, Apn1, Apn2, and Rad1, does (Figures 1 and 4). This important difference allows us to propose that it is the hyperactivation of Apn1/2 that is most crucial for the generation of DSBs at closely juxtaposed sites of Zeocin-induced damage. Apn1/2 appears to be misregulated by nuclear localized G-actin.

Here we present evidence that Apn1/APE1 binds actin both in yeast extracts and in vitro (Figure 5). While our data do not resolve whether or not actin plays a physiological role in the repair process or in the generation of DSBs, our data suggest that depolymerized actin either activates Apn1/2 activity, or increases its abundance in the nucleus, or facilitates access to the second lesion, allowing premature processing of the second site and driving DSB formation (Figure 6). This assumes that two adjacent DNA lesions are processed by Apn1/2 under normal conditions in a sequential manner, minimizing the chance of spontaneous DSB formation.

The AP endonucleases, encoded by *APN1* and *APN2* in yeast and APE1 in mammals, have complex roles in the response to DNA damage. The mammalian APE1 enzyme not only cuts at the apurinic/apyrimidinic sites generated by glycosylase-mediated excision of damaged bases, but together with Thymine DNA glycosylase (TDG), APE1 catalyses the first steps in BER-dependent active demethylation of cytosines (reviewed in ^3^). In this context, APE1 was also shown to trigger the release of TDG and other glycosylases from abasic sites. This release of glycosylase may trigger premature processing.

The mammalian APE1, like yeast Apn1/2, also participates in what has recently been called the BER/NIR switch ^52, 53, 57, 58^. In the NIR pathway, APE1 (or Apn1/2) bypasses the action of the glycosylase and generates a nick that retains the 5’ damaged nucleotide and generates a 3’OH group. In this function AP enzymes can recognize and process a diverse set of lesions, including oxidized pyrimidines, formamido-pyrimidines, exocyclic DNA bases and uracil, bulky lesions and UV-induced 6-4 photoproducts (reviewed in ^3^). The switch from a glycosylase-dependent to a glycosylase-independent cleavage mode appears to depend on allosteric changes in APE1 structure, possibly controlled by its N-terminal tail ^57^. We hypothesize that the fragmentation that we see following Zeocin treatment in G1 phase cells may arise from Apn1/2 acting in a glycosylase-independent manner, since deletion of Ogg1 and Ntg1/2 did not block the massive fragmentation of the genome in response to Zeocin in G1 phase.

We propose that actin depolymerization either by Tor2 inhibition, LatA treatment, or Las17 degradation, directly promotes simultaneous processing of adjacent Zeocin-induced lesions. We note that in S/G2 phase, the defect could also arise from hyperactivity of DNA polymerases, the loss of which reduces YCS in S-phase cells only (Figure 3). It is possible that strand hyperextension by DNA Polδ could also generate DSBs if the elongating strand encounters a second lesion that has already been processed to generate a ss nick. DNA Polδ indeed is able to fill gaps and efficiently catalyze subsequent strand displacement synthesis ^42^. This might also be promoted by the helicase activity of Dnase2 (*DNA2*) whose loss also confers partial resistance the in presence of Zeocin and TORC2 inhibition (data not shown). On the other hand, since Apn1 and Apn2 both possess 3’ to 5’ exonuclease activity ^51, 59^, the conversion of two ss nicks to a DSB could be mediated by the AP endonuclease extending the gap on one strand such that it encounters a nick on the other strand, generating a DSB on its own (Figure 6).

The effects of actin depolymerization on Apn1 activity appears not to be cell-cycle dependent. Thus, one mechanism affecting this might entail the release of the mitochondrial pool of Apn1. Apn1 not only repairs oxidized bases in the nucleus, but it serves the same purpose in mitochondria ^60^. Given that actin depolymerization appears to trigger mitochondrial reticularization ^61, 62^, the mitochondrial complement of Apn1 may become mistargeted to the nucleus under conditions of actin depolymerization, enhancing the Apn1 nuclear pool. This is consistent with our finding that overexpression of Apn1 with a nuclear localization signal renders yeast hypersensitive to Zeocin (Suppl. Figure 4). In an alternative model, the nuclear pool of Apn1/2 might not change, but rather a co-factor that coordinates the sequential processing of adjacent lesions by Apn1/2, may respond to nuclear actin pools. We note that Apn1 and APE1 both bind actin (Figure 5). However, we were unable to monitor a significant stimulation of APE1 nuclease activity by actin *in vitro* (data not shown). Instead, the accentuation of abasic site processing may stem from enhanced glycosylase eviction, nucleosome eviction or enhanced DNA polymerase elongation rates, which might all render the second nearby lesion more accessible. Such events might also be influenced by nucleosome remodelers, such as INO80 or SWR1, both of which contain an essential actin-Arp4 dimer ^21^. How an excess of nuclear G-actin influences INO80 remodeler activity, however, is unknown.

The induction of DSBs by non-canonical BER processing occurs naturally in B lymphocytes during antibody class switch recombination (CSR; reviewed in ^2^). For this process, cytosine deamination by the Activation Induced Deaminase (AID) enzyme, generates uracil lesions within the switch regions of the immunoglobulin heavy chain locus. The switch regions are highly repetitive sequences that are rich in CpGs; thus, uracils can be generated in close proximity and on opposite strands. Further processing by the uracil DNA glycosylase (UNG) and APE1, generates ssDNA nicks, and when in close proximity, this results in DSBs ^63, 64^. Such breaks can also be generated by uncoordinated processing by the BER machinery and the mismatch repair machinery (MMR), acting on opposite strands (reviewed in ^65^). For CSR, further repair by NHEJ generates a translocation between two different switch regions, both containing DSBs, which alters the constant region associated with the expressed antibody. There are several parallels between YCS and CSR, most notably the role of non-canonical BER in generating DSBs. It is interesting to note that a hyperactive and mistargeted form of AID is highly toxic as its lesions are misprocessed to generate excessive DSBs, a phenotype reminiscent of YCS ^66^.

Beyond this rather specialized case of APE1-generated breaks in antibody class switch recombination, we have examined other ways in which this phenomenon might be relevant for human disease. As mentioned above, several papers have argued for a role of actin and actin filament nucleating proteins in DSB repair ^26, 27, 67^. The lethality we observe following F-actin misregulation does not depend on nor mimic loss of DSB repair, whether by HR, NHEJ, or both. In fact, we see no link between the effect of actin depolymerization and DSB repair in yeast. Moreover, in primary human fibroblasts and HCT111 cells, we find a synergistic lethality provoked by actin depolymerization in the presence of Zeocin (Hurst *et al*., in revision). It is unknown at present whether this synergy requires APE1. Intriguingly we find that LatA treatment in mammalian cells decreases the intensity of GFP-XRCC1 and GFP-PCNA recruitment to laser-induced damage (VH and SMG, unpublished). This damage is likely to include thymidine dimers, adducts and other oxidization damage that might serve as substrates for NIR.

A recent report finds that low dose doxorubicin, a topoisomerase II inhibitor, can be potentiated by combining it with subtoxic levels of actin modifying substances – notably jasplakinolide, Condramide B, or Latrunculin B ^68^. These authors present evidence, consistent with ours, that DNA repair is less efficient following actin depolymerization. In a separate study, however, it was proposed that actin depolymerization increases Doxorubicin uptake, thus generating more damage, rather than inhibiting repair ^69^. We conclude that in mammalian cells, the exact mode of synergism between G-actin and modified base repair, is unclear. While it is too early to advocate combining actin depolymerizing agents with DNA damaging agents in combinatorial cancer therapies, this approach merits testing ^68, 70^. Our current study clearly illustrates synergistic lethality between low level Zeocin-induced damage and TORC2 inhibition in yeast. The most immediate application of our findings may be, therefore, the development of highly effective anti-fungal treatments.

## Acknowledgements

We thank H. Araki, K. Shirahige, and M. Kanemaki for the generous gifts of yeast strains, and S.B. Helliwell (NIBR) for the TOR inhibitors. This work was supported by the Human Frontiers Science Program Grant: Actin and Actin-related proteins. We also acknowledge the generous financial support of the Swiss National Science Foundation grant number 31003A-176286 to S.M.G for support of V.H. and K.S. and 31003A-149989 to J.J. for support of S.B. C.G. was supported by a FP7 Marie-Curie Intra-European Fellowship. We thank all Gasser laboratory members for extensive discussions over the years and the Novartis Research Foundation for continued support. Finally, we are very grateful to Sam Wilson, Primo Schaer and Ulrich Hübscher for valuable discussions.

## Competing interests

The authors declare that no competing interests exist.

## Materials and Methods

### Cell culture, strains, plasmids and chemicals

Yeast strains and plasmids used in this study are listed in the Supplementary yeast and plasmid table 1. Yeast cells were cultured in SC (synthetic complete-2% glucose) medium at 30°C unless otherwise indicated. To deplete auxin dependent degron target, 0.5 mM indoleacetic acid (IAA, Sigma Aldrich) was added to the culture. The TORC1 and TORC2 inhibitors CMB4563 and NHP-BHS345 are closely related imidazoquinolines, with CMB being roughly 10-fold more effective in the YCS assay (Hurst *et al*., in revision). The compounds are dissolved in DMSO and were the kind gift of Stephen B. Helliwell (Novartis Institutes of Biomedical Research, Basel, Switzerland). In mammalian cells they indiscriminately repress a broad range of PIK Kinases, while in yeast particularly CMB4563 preferentially inhibits TORC2.

### CHEF gel analysis

Yeast genomic DNA was prepared in an agarose plug as described in the Instruction Manual (Bio-Rad, CHEF-DR II) with slight modifications. Yeast cells were span and washed with ice-cold 50 mM EDTA-NaOH (pH8.0), then cell pellet was suspended in Zymolyase buffer (50 mM Na-phosphate pH7.0, 50 mM EDTA, 1 mM DTT) and embedded in 1% agarose plug. Genomic DNA were prepared in the agarose plug by 0.4 mg/ml Zymolyase (20T, Seikagaku) treatment in Zymolyase buffer at 37°C for 1h, followed by 1 mg/ml Proteinase K digestion in 10 mM Tris-HCl pH7.5, 50 mM EDTA, 1% sodium N-lauroylsarcosinate at 50°C more than 12h, except genomic DNA preparation in Figure 1 was performed at 30°C. Chromosomal DNA was separated in 1% agarose/0.5 × TBE on the CHEF-DR II Pulsed Field Electrophoresis Systems (Bio-Rad) as follows: 14°C, 6 V /cm, 60 s switch time for 15 h, then 90 s switch time for 7 h. Chromosomal DNA was stained with ethidium bromide or SYBR safe, and imaged on the Chem Doc XRS system (Bio-Rad) or Typhoon FLA 9500 scanner (GE Healthcare). Signal intensity of each lane was measured by Image J program. In the Image J program, a region of interest (ROI) spanning from the largest chromosome band to ~20 kb in size was created and the signal intensity was measured using line plot profiling. The same ROI was applied for each lane in a CHEF gel image. Signal intensity was divided to region A above 0.57 Mb which covers the largest chromosome 4 and 12 band to chromosome 5 and 8, and region B below 0.57 Mbp which covers a majority of DSB signal and small chromosome 1, 3, 6, and 9. A signal ratio B over A was calculated and indicated under the each CHEF gel image, representing the degree of chromosome fragmentation.

Quantification of the mean number of breaks per chromosome was determined based on the assumption that DSBs are independently and uniformly distributed, i.e. they occur at the same frequency (*λ*) per unit length of DNA in all the chromosomes. Thus the number of breaks in chromosome i follows a Poisson distribution of parameter *λS*_*i*_, where *S*_*i*_ is the size of chromosome *i*. In this model, the mean fragment size over all the chromosomes is 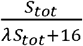, where *S*_*tot*_ is the length of the genome. Hence the mean number of breaks per chromosome can be calculated as a function of the mean fragment size. Given the estimated mean fragment size in the YCS experiments is the (i.e. maximal intensity distribution curve 100±20 kb). The corresponding mean number of breaks per chromosome are from the smallest to largest: 1.9595; 6.9213; 2.6949; 13.0389; 4.9100; 2.2994; 9.2854; 4.7889; 3.7441; 6.3474; 5.6755; 9.1768; 7.8682; 6.6758; 9.2884; 8.0694 (see Figure 1C-D).

### In vitro plasmid nicking assay

pRichi, pGEM13Zf(+) derivative supercoiled plasmid^71^ was incubated with Zeocin (Life Technologies) in MMR buffer (20 mM Tris-HCl ph 7.6, 40 mM KCl, 5 mM MgCl_2_, 50 ng/μL BSA, 1 mM Glutathione) for 60 min at 37°C. The sample was divided in two, then 0.002 U of recombinant hOGG1 (Trevigen, 4130-100-E) was added to one sample and further incubated for 30 min at 37°C. Plasmid DNA was purified by NucleoSpin column (Macherey-Nagel), then analysed on a 1% agarose gel containing RedSafe™ (Sigma-Aldrich) in 1 × TAE.

### Western blot and antibody

Protein extracts were separated by SDS-PAGE or NuPAGE (Invitrogen) and transferred to nitrocellulose membranes BA-85 (Whatman) for probing. Antibodies used were: anti-actin (Millipore, Mab1501 clone C4), anti-MCM2 (Santa Cruz, SC-6680), anti-tubulin (Thermo Fisher Scientific: MA1-80017), and anti-MYC (9E10).

### Actin-bicyclic peptide and pull-down

Detail information of actin bicyclic peptides including identification and characterization are described in Gübeli *et al*. ^56^. 10 μg of Anti-fluorescein (FITC) mouse antibody (Jackson Immuno Res., clone IF8-IE4) was pre-coupled with 50 μl of Protein G-Dynal beads (Thermo Fischer Scientific), then 2 nmol of FITC labeled bicyclic peptides A18 (sequence: ACREGQVACMVRKFECG), A15 (sequence: ACYRQWNKCENGWVRCG), and TATA2 (sequence: ANCPLVCAPRCR) were bound to anti-FITC-coupled beads in the lysis buffer (50 mM HEPES pH7.5, 20 mM NaCl, 1 mM EDTA, 0.1% Triton X-100, protease inhibitor cocktail (Roche)) for 1.5h at room temperature, then washed three times with lysis buffer before use. Exponentially growing yeast cells (~1 × 10^7^ cells/ml, 200 ml) were pelleted and washed once with ice-cold PBS. Cell pellet was resuspended in 0.8 ml lysis buffer and the cell lysate was prepared by beat beating with Zerconia beads, 6.5 Hz, 60 sec, 4 times at 4°C. Cell lysate was clarified by centrifugation 12000 × g, 5 min at 4°C. 150 μl of total cell lysate was incubated with bicyclic peptide coupled Dynal beads (50 μl) for 1.5h at 4°C with constant rotation, then Dynal beads were washed three times with excess lysis buffer containing 0.1% TritonX-100 and 20 mM salt at 4°C. Pull-down proteins were eluted from the beads with 60 μl of 100 mM Glycine pH2.0, 0.1% Triton X100, then pH was quickly neutralized by 3.5 μl of 1M Tris-HCl pH 9.5. Protein sample was denatured and boiled in 1x NuPAGE sample buffer, and analysed by NuPAGE followed by silver staining and Western blot.

---

**Figure 1 Suppl. 1.**
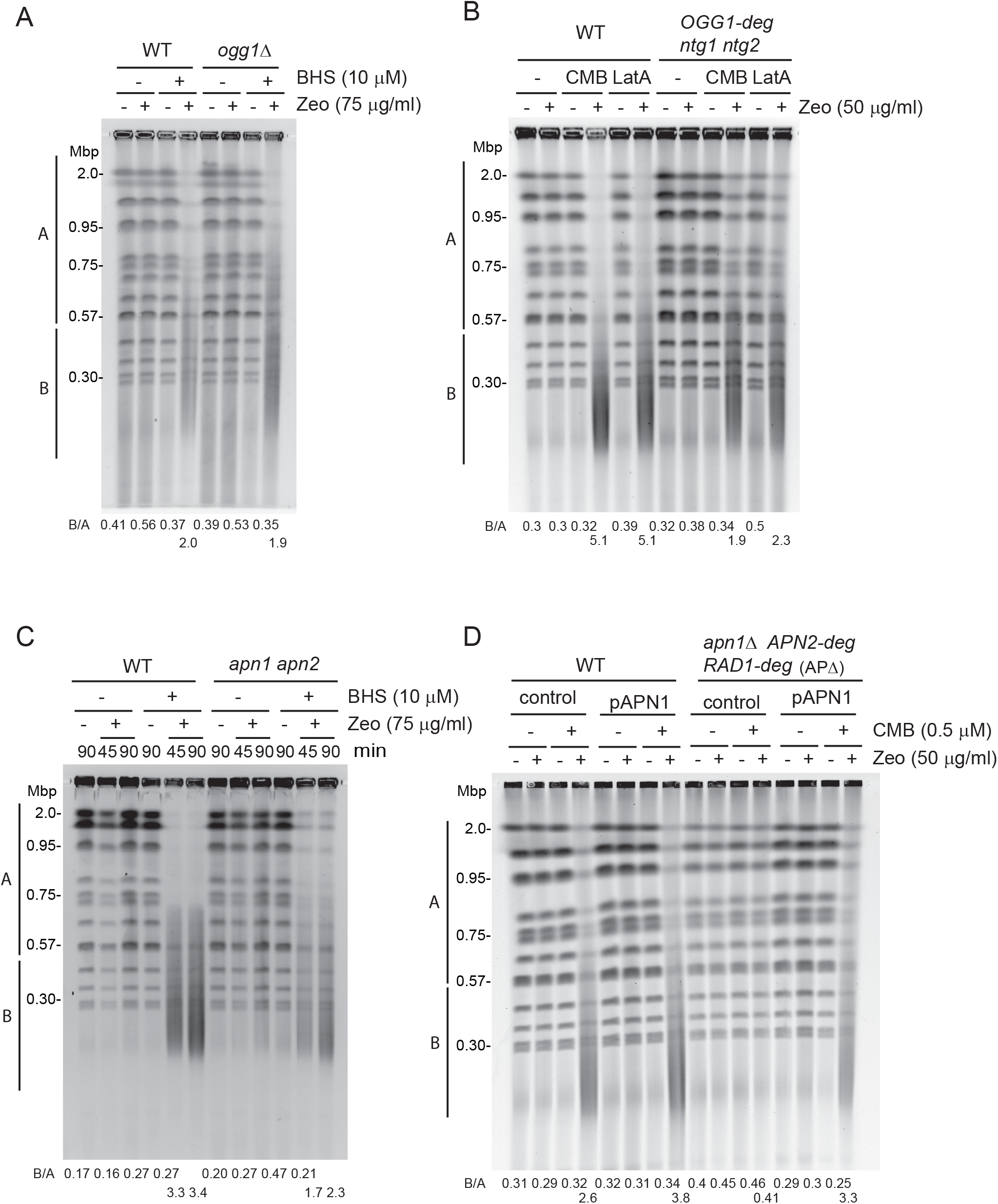

**Figure 2 Suppl. 1.**
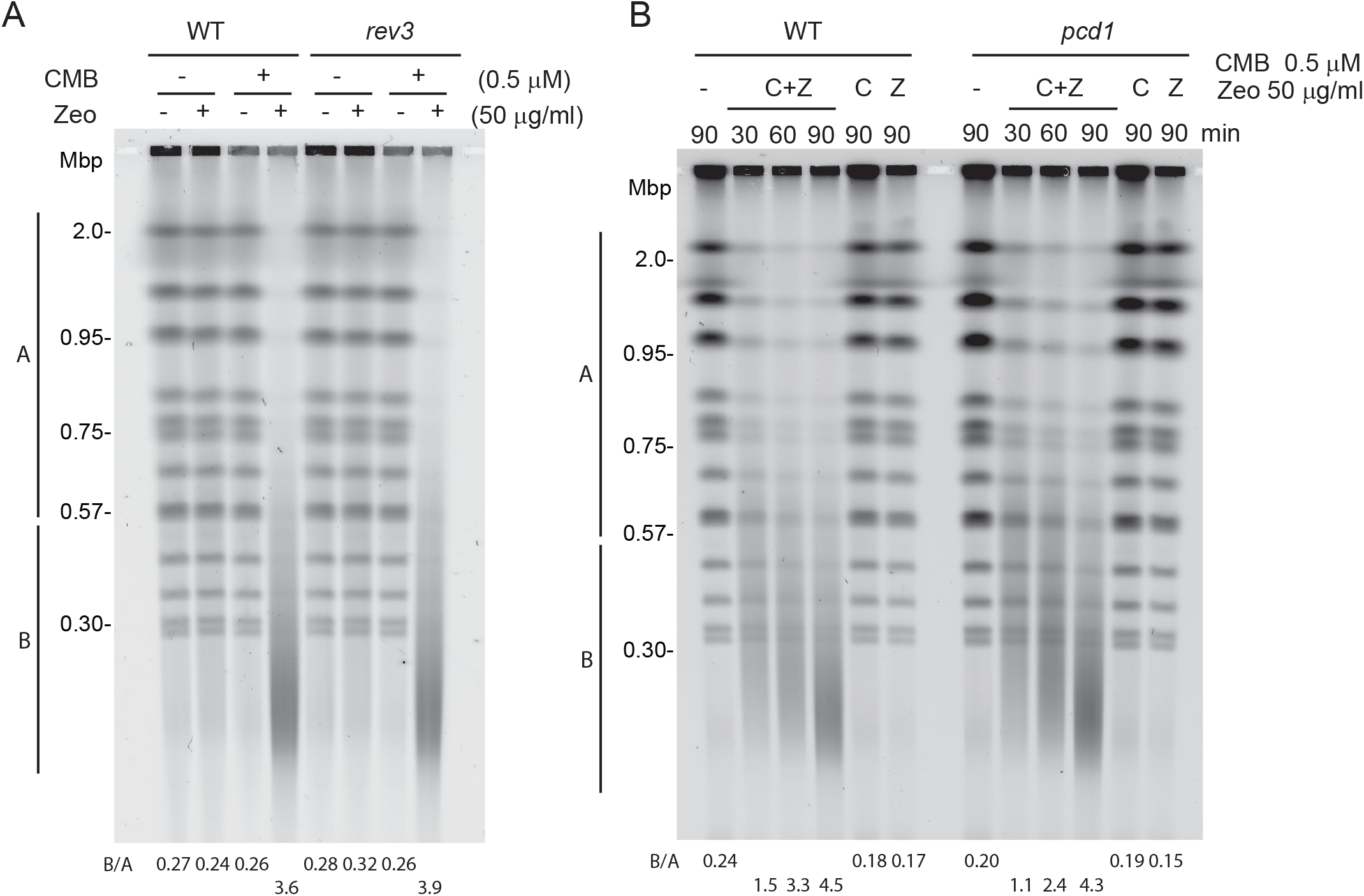

**Figure 3 Suppl. 1.**
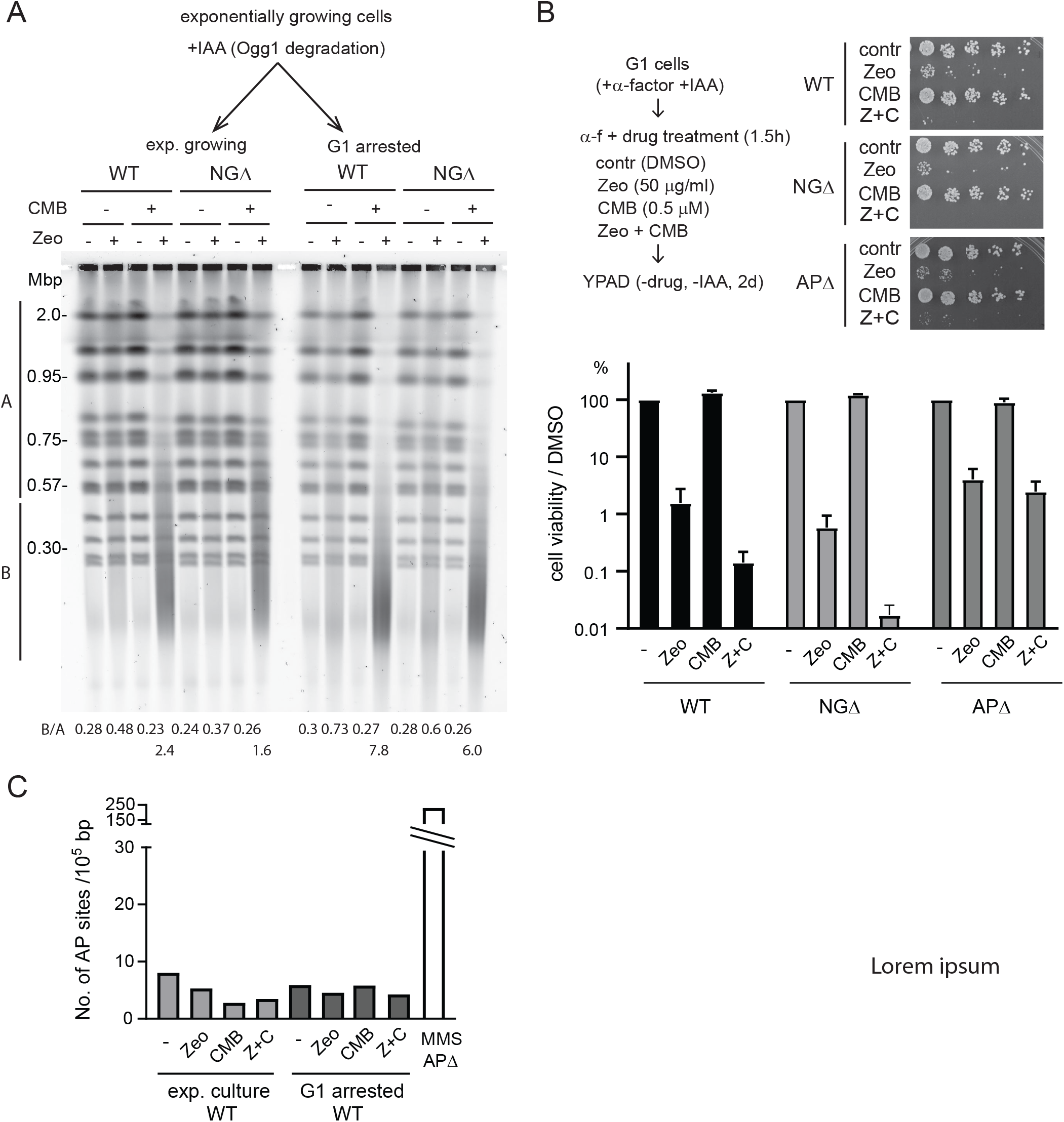

**Figure 4 Suppl. 1.**
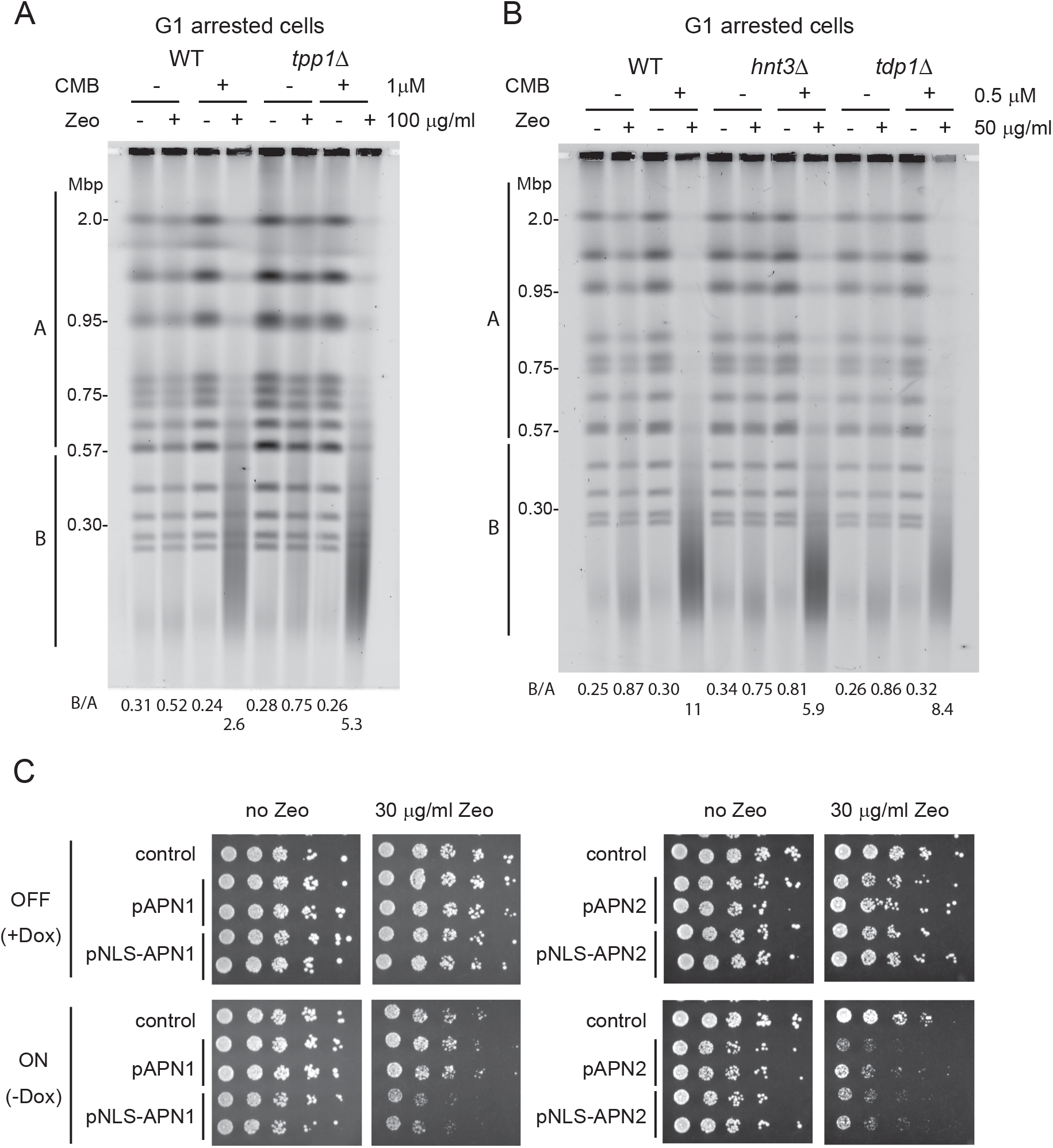

## Notes

### Competing Interest Statement

The authors have declared no competing interest.

## References

1. Lindahl T, Barnes DE. Repair of endogenous DNA damage. Cold Spring Harb Symp Quant Biol 65, 127–133 (2000).

2. Methot SP, Di Noia JM. Molecular Mechanisms of Somatic Hypermutation and Class Switch Recombination. Adv Immunol 133, 37–87 (2017).

3. Popov AV, et al. Reading Targeted DNA Damage in the Active Demethylation Pathway: Role of Accessory Domains of Eukaryotic AP Endonucleases and Thymine-DNA Glycosylases. J Mol Biol, (2019).

4. van Loon B, Markkanen E, Hubscher U. Oxygen as a friend and enemy: How to combat the mutational potential of 8-oxo-guanine. DNA Repair (Amst) 9, 604–616 (2010).

5. de la Torre-Ruiz MA, Pujol N, Sundaran V. Coping with oxidative stress. The yeast model. Curr Drug Targets 16, 2–12 (2015).

6. Boiteux S, Jinks-Robertson S. DNA repair mechanisms and the bypass of DNA damage in Saccharomyces cerevisiae. Genetics 193, 1025–1064 (2013).

7. Meadows KL, Song B, Doetsch PW. Characterization of AP lyase activities of Saccharomyces cerevisiae Ntg1p and Ntg2p: implications for biological function. Nucleic acids research 31, 5560–5567 (2003).

8. Girard PM, Guibourt N, Boiteux S. The Ogg1 protein of Saccharomyces cerevisiae: a 7,8-dihydro-8-oxoguanine DNA glycosylase/AP lyase whose lysine 241 is a critical residue for catalytic activity. Nucleic acids research 25, 3204–3211 (1997).

9. Dalhus B, et al. Separation-of-function mutants unravel the dual-reaction mode of human 8-oxoguanine DNA glycosylase. Structure 19, 117–127 (2011).

10. Wang Z, Wu X, Friedberg EC. DNA repair synthesis during base excision repair in vitro is catalyzed by DNA polymerase epsilon and is influenced by DNA polymerases alpha and delta in Saccharomyces cerevisiae. Molecular and cellular biology 13, 1051–1058 (1993).

11. Blank A, Kim B, Loeb LA. DNA polymerase delta is required for base excision repair of DNA methylation damage in Saccharomyces cerevisiae. Proc Natl Acad Sci U S A 91, 9047–9051 (1994).

12. Gellon L, Carson DR, Carson JP, Demple B. Intrinsic 5’-deoxyribose-5-phosphate lyase activity in Saccharomyces cerevisiae Trf4 protein with a possible role in base excision DNA repair. DNA Repair (Amst) 7, 187–198 (2008).

13. Almeida KH, Sobol RW. A unified view of base excision repair: lesion-dependent protein complexes regulated by post-translational modification. DNA Repair (Amst) 6, 695–711 (2007).

14. Chalissery J, Jalal D, Al-Natour Z, Hassan AH. Repair of Oxidative DNA Damage in Saccharomyces cerevisiae. DNA Repair (Amst) 51, 2–13 (2017).

15. Saleh-Gohari N, Bryant HE, Schultz N, Parker KM, Cassel TN, Helleday T. Spontaneous homologous recombination is induced by collapsed replication forks that are caused by endogenous DNA single-strand breaks. Molecular and cellular biology 25, 7158–7169 (2005).

16. Kuzminov A. Single-strand interruptions in replicating chromosomes cause double-strand breaks. Proc Natl Acad Sci U S A 98, 8241–8246 (2001).

17. Ma W, Panduri V, Sterling JF, Van Houten B, Gordenin DA, Resnick MA. The transition of closely opposed lesions to double-strand breaks during long-patch base excision repair is prevented by the coordinated action of DNA polymerase delta and Rad27/Fen1. Molecular and cellular biology 29, 1212–1221 (2009).

18. Shimada K, et al. TORC2 signaling pathway guarantees genome stability in the face of DNA strand breaks. Mol Cell 51, 829–839 (2013).

19. Povirk LF. DNA damage and mutagenesis by radiomimetic DNA-cleaving agents: bleomycin, neocarzinostatin and other enediynes. Mutat Res 355, 71–89 (1996).

20. Roelants FM, et al. TOR Complex 2-Regulated Protein Kinase Fpk1 Stimulates Endocytosis via Inhibition of Ark1/Prk1-Related Protein Kinase Akl1 in Saccharomyces cerevisiae. Molecular and cellular biology 37, (2017).

21. Seeber A, Hauer M, Gasser SM. Nucleosome remodelers in double-strand break repair. Curr Opin Genet Dev 23, 174–184 (2013).

22. Dion V, Shimada K, Gasser SM. Actin-related proteins in the nucleus: life beyond chromatin remodelers. Curr Opin Cell Biol 22, 383–391 (2010).

23. Kapoor P, Shen X. Mechanisms of nuclear actin in chromatin-remodeling complexes. Trends Cell Biol 24, 238–246 (2014).

24. Hurst V, Shimada K, Gasser SM. Nuclear Actin and Actin-Binding Proteins in DNA Repair. Trends Cell Biol 29, 462–476 (2019).

25. Belin BJ, Lee T, Mullins RD. DNA damage induces nuclear actin filament assembly by Formin - 2 and Spire-(1/2) that promotes efficient DNA repair. [corrected]. Elife 4, e07735 (2015).

26. Schrank BR, et al. Nuclear ARP2/3 drives DNA break clustering for homology-directed repair. Nature 559, 61–66 (2018).

27. Caridi CP, et al. Nuclear F-actin and myosins drive relocalization of heterochromatic breaks. Nature 559, 54–60 (2018).

28. Avendano C, Menéndez JC. Medicinal Chemistry of Anticancer Drugs. Elsevier Science (2008).

29. Fu D, Calvo JA, Samson LD. Balancing repair and tolerance of DNA damage caused by alkylating agents. Nat Rev Cancer 12, 104–120 (2012).

30. Lundin C, et al. Methyl methanesulfonate (MMS) produces heat-labile DNA damage but no detectable in vivo DNA double-strand breaks. Nucleic acids research 33, 3799–3811 (2005).

31. Bourgoint C, et al. Target of rapamycin complex 2-dependent phosphorylation of the coat protein Pan1 by Akl1 controls endocytosis dynamics in Saccharomyces cerevisiae. J Biol Chem 293, 12043–12053 (2018).

32. Chakrabarti S, Makrigiorgos GM, O’Brien K, Bump E, Kassis AI. Measurement of hydroxyl radicals catalyzed in the immediate vicinity of DNA by metal-bleomycin complexes. Free Radic Biol Med 20, 777–783 (1996).

33. Boiteux S, Radicella JP. The human OGG1 gene: structure, functions, and its implication in the process of carcinogenesis. Arch Biochem Biophys 377, 1–8 (2000).

34. Guillet M, Boiteux S. Endogenous DNA abasic sites cause cell death in the absence of Apn1, Apn2 and Rad1/Rad10 in Saccharomyces cerevisiae. EMBO J 21, 2833–2841 (2002).

35. Kelley MR, Kow YW, Wilson DM, 3rd. Disparity between DNA base excision repair in yeast and mammals: translational implications. Cancer Res 63, 549–554 (2003).

36. Prindle MJ, Loeb LA. DNA polymerase delta in DNA replication and genome maintenance. Environ Mol Mutagen 53, 666–682 (2012).

37. Auerbach PA, Demple B. Roles of Rev1, Pol zeta, Pol32 and Pol eta in the bypass of chromosomal abasic sites in Saccharomyces cerevisiae. Mutagenesis 25, 63–69 (2010).

38. Tseng HM, Tomkinson AE. Processing and joining of DNA ends coordinated by interactions among Dnl4/Lif1, Pol4, and FEN-1. J Biol Chem 279, 47580–47588 (2004).

39. Lemos BR, et al. CRISPR/Cas9 cleavages in budding yeast reveal templated insertions and strand-specific insertion/deletion profiles. Proc Natl Acad Sci U S A 115, E2040–E2047 (2018).

40. Johnson RE, Prakash L, Prakash S. Pol31 and Pol32 subunits of yeast DNA polymerase delta are also essential subunits of DNA polymerase zeta. Proc Natl Acad Sci U S A 109, 12455–12460 (2012).

41. Sukhanova MV, D’Herin C, van der Kemp PA, Koval VV, Boiteux S, Lavrik OI. Ddc1 checkpoint protein and DNA polymerase varepsilon interact with nick-containing DNA repair intermediate in cell free extracts of Saccharomyces cerevisiae. DNA Repair (Amst) 10, 815–825 (2011).

42. Ganai RA, Zhang XP, Heyer WD, Johansson E. Strand displacement synthesis by yeast DNA polymerase epsilon. Nucleic acids research 44, 8229–8240 (2016).

43. Freudenthal BD, et al. Uncovering the polymerase-induced cytotoxicity of an oxidized nucleotide. Nature 517, 635–639 (2015).

44. Caglayan M, Horton JK, Dai DP, Stefanick DF, Wilson SH. Oxidized nucleotide insertion by pol beta confounds ligation during base excision repair. Nat Commun 8, 14045 (2017).

45. Fujikawa K, Kamiya H, Yakushiji H, Fujii Y, Nakabeppu Y, Kasai H. The oxidized forms of dATP are substrates for the human MutT homologue, the hMTH1 protein. J Biol Chem 274, 18201–18205 (1999).

46. Xiao W, Samson L. In vivo evidence for endogenous DNA alkylation damage as a source of spontaneous mutation in eukaryotic cells. Proc Natl Acad Sci U S A 90, 2117–2121 (1993).

47. Ma W, Resnick MA, Gordenin DA. Apn1 and Apn2 endonucleases prevent accumulation of repair-associated DNA breaks in budding yeast as revealed by direct chromosomal analysis. Nucleic acids research 36, 1836–1846 (2008).

48. Vance JR, Wilson TE. Repair of DNA strand breaks by the overlapping functions of lesion-specific and non-lesion-specific DNA 3’ phosphatases. Molecular and cellular biology 21, 7191–7198 (2001).

49. Daley JM, Wilson TE, Ramotar D. Genetic interactions between HNT3/Aprataxin and RAD27/FEN1 suggest parallel pathways for 5’ end processing during base excision repair. DNA Repair (Amst) 9, 690–699 (2010).

50. El-Khamisy SF, Katyal S, Patel P, Ju L, McKinnon PJ, Caldecott KW. Synergistic decrease of DNA single-strand break repair rates in mouse neural cells lacking both Tdp1 and aprataxin. DNA Repair (Amst) 8, 760–766 (2009).

51. Ishchenko AA, Yang X, Ramotar D, Saparbaev M. The 3’->5’ exonuclease of Apn1 provides an alternative pathway to repair 7,8-dihydro-8-oxodeoxyguanosine in Saccharomyces cerevisiae. Molecular and cellular biology 25, 6380–6390 (2005).

52. Ischenko AA, Saparbaev MK. Alternative nucleotide incision repair pathway for oxidative DNA damage. Nature 415, 183–187 (2002).

53. Dyakonova ES, Koval VV, Lomzov AA, Ishchenko AA, Fedorova OS. Apurinic/apyrimidinic endonuclease Apn1 from Saccharomyces cerevisiae is recruited to the nucleotide incision repair pathway: Kinetic and structural features. Biochimie 152, 53–62 (2018).

54. Brault V, Sauder U, Reedy MC, Aebi U, Schoenenberger CA. Differential epitope tagging of actin in transformed Drosophila produces distinct effects on myofibril assembly and function of the indirect flight muscle. Mol Biol Cell 10, 135–149 (1999).

55. Melak M, Plessner M, Grosse R. Actin visualization at a glance. J Cell Sci 130, 525–530 (2017).

56. Gübeli R, et al. In Vitro-Evolved Peptides Mimic a Binding Motif of the G-Actin-Binding Protein Thymosin-B4 and Serve as Research Tools. ChemRxiv Preprint, (2020).

57. Gros L, Ishchenko AA, Ide H, Elder RH, Saparbaev MK. The major human AP endonuclease (Ape1) is involved in the nucleotide incision repair pathway. Nucleic acids research 32, 73–81 (2004).

58. Ishchenko AA, Sanz G, Privezentzev CV, Maksimenko AV, Saparbaev M. Characterisation of new substrate specificities of Escherichia coli and Saccharomyces cerevisiae AP endonucleases. Nucleic acids research 31, 6344–6353 (2003).

59. Unk I, Haracska L, Prakash S, Prakash L. 3’-phosphodiesterase and 3’-->5’ exonuclease activities of yeast Apn2 protein and requirement of these activities for repair of oxidative DNA damage. Molecular and cellular biology 21, 1656–1661 (2001).

60. Acevedo-Torres K, Fonseca-Williams S, Ayala-Torres S, Torres-Ramos CA. Requirement of the Saccharomyces cerevisiae APN1 gene for the repair of mitochondrial DNA alkylation damage. Environ Mol Mutagen 50, 317–327 (2009).

61. Altmann K, Westermann B. Role of essential genes in mitochondrial morphogenesis in Saccharomyces cerevisiae. Mol Biol Cell 16, 5410–5417 (2005).

62. Boldogh IR, Pon LA. Interactions of mitochondria with the actin cytoskeleton. Biochim Biophys Acta 1763, 450–462 (2006).

63. Masani S, Han L, Yu K. Apurinic/apyrimidinic endonuclease 1 is the essential nuclease during immunoglobulin class switch recombination. Molecular and cellular biology 33, 1468–1473 (2013).

64. Rada C, Williams GT, Nilsen H, Barnes DE, Lindahl T, Neuberger MS. Immunoglobulin isotype switching is inhibited and somatic hypermutation perturbed in UNG-deficient mice. Curr Biol 12, 1748–1755 (2002).

65. Zanotti KJ, Gearhart PJ. Antibody diversification caused by disrupted mismatch repair and promiscuous DNA polymerases. DNA Repair (Amst) 38, 110–116 (2016).

66. Zahn A, et al. Activation induced deaminase C-terminal domain links DNA breaks to end protection and repair during class switch recombination. Proc Natl Acad Sci U S A 111, E988–997 (2014).

67. Andrin C, et al. A requirement for polymerized actin in DNA double-strand break repair. Nucleus 3, 384–395 (2012).

68. Pfitzer L, et al. Targeting actin inhibits repair of doxorubicin-induced DNA damage: a novel therapeutic approach for combination therapy. Cell Death Dis 10, 302 (2019).

69. Takeshita H, Kusuzaki K, Ashihara T, Gebhardt MC, Mankin HJ, Hirasawa Y. Actin organization associated with the expression of multidrug resistant phenotype in osteosarcoma cells and the effect of actin depolymerization on drug resistance. Cancer Lett 126, 75–81 (1998).

70. Weisman R, Cohen A, Gasser SM. TORC2-a new player in genome stability. EMBO Mol Med 6, 995–1002 (2014).

71. Fischer F, Baerenfaller K, Jiricny J. 5-Fluorouracil is efficiently removed from DNA by the base excision and mismatch repair systems. Gastroenterology 133, 1858–1868 (2007).

